# Investigating the consequences of chronic short sleep for metabolism and survival of oxidative stress

**DOI:** 10.1101/2024.12.01.626207

**Authors:** Samantha J. Tener, Chloe E. Kim, Jungwon Lee, Kairaluchi Oraedu, Jared A. Gatto, Timothy Y. Chang, Carly Lam, Ryan Schanta, Meaghan S. Jankowski, Scarlet J. Park, Jennifer M. Hurley, Matthew Ulgherait, Julie C. Canman, William W. Ja, Douglas B. Collins, Mimi Shirasu-Hiza

## Abstract

In previous work, we found that short sleep caused sensitivity to oxidative stress; here we set out to characterize the physiological state of a diverse group of chronically short-sleeping mutants during hyperoxia as an acute oxidative stress. Using RNA-sequencing analysis, we found that short-sleeping mutants had a normal transcriptional oxidative stress response relative to controls. In both short-sleeping mutants and controls, hyperoxia led to downregulation of glycolytic genes and upregulation of genes involved in fatty acid metabolism, reminiscent of metabolic shifts during sleep. We hypothesized that short-sleeping mutants may be sensitive to hyperoxia because of defects in metabolism. Consistent with this, short-sleeping mutants were sensitive to starvation. Using metabolomics, we identified a pattern of low levels of long chain fatty acids and lysophospholipids in short-sleeping mutants relative to controls during hyperoxia, suggesting a defect in lipid metabolism. Though short-sleeping mutants did not have common defects in many aspects of lipid metabolism (basal fat stores, usage kinetics during hyperoxia, respiration rates, and cuticular hydrocarbon profiles), they were all sensitive to dehydration, suggesting a general defect in cuticular hydrocarbons, which protect against dehydration. To test the bi-directionality of sleep and lipid metabolism, we tested flies with both diet-induced obesity and genetic obesity. Flies with diet-induced obesity had no sleep or oxidative stress phenotype; in contrast, the lipid metabolic mutant, *brummer*, slept significantly more than controls but was sensitive to oxidative stress. Previously, all short sleepers tested were sensitive and all long sleepers resistant to oxidative stress. *brummer* mutants, the first exceptions to this rule, lack a key enzyme required to mobilize fat stores, suggesting that a defect in accessing lipid stores can cause sensitivity to oxidative stress. Taken together, we found that short-sleeping mutants have many phenotypes in common: sensitivity to oxidative stress, starvation, dehydration, and defects in lipid metabolites. These results argue against a specific role for sleep as an antioxidant and suggest the possibility that lipid metabolic defects underlie the sensitivity to oxidative stress of short-sleeping mutants.

## INTRODUCTION

Sleep is a widespread, evolutionarily conserved behavior of animals.^1–5^ The diverse range of organisms that exhibit sleep behavior is often seen as evidence for sleep’s fundamental importance in organismal well-being. Moreover, lack of sleep is strongly associated with disease. Chronic short-sleep has been associated with the development of metabolic disorders^6–8^, particularly obesity and type 2 diabetes^9^, coronary heart disease and stroke^10^, dementia, Alzheimer’s disease and cognitive decline^11–15^, Parkinsonian disorders^16^, kidney disease^17^, liver disease^18,19^, particularly non-alcoholic fatty liver disease, gastrointestinal diseases^20^, immune dysfunction^21^, and overall mortality^22–24^. Thus, a lack of sleep is associated with multiple pathological consequences throughout the body. Because of the observed association with such a wide variety of diseases across many different organs and tissues, sleep is hypothesized to promote health through an essential and ubiquitous cellular process.

Using *Drosophila melanogaster*, an established model system for the study of sleep^25,26^, our lab previously published work demonstrating that sleep functions to promote oxidative stress defense^27^, a theory first proposed by Eric Reimund^28^. Oxidative stress is caused by reactive oxygen species (ROS), which are metabolic byproducts generated by mitochondria. ROS cause damage to macromolecules essential for a cell’s function and oxidative stress occurs when toxic ROS accumulation and damage outweigh a cell’s antioxidant response. Other groups have also shown that markers of oxidative stress increase as a consequence of sleep loss.^29–35^ However, in most of these studies, rodents were subjected to acute sleep deprivation, which does not mirror the chronic sleep loss linked to disease in humans. Furthermore, these studies employed sleep deprivation protocols that were themselves stress-inducing, which may have confounded the measurements of stress biomarkers. Thus, genetically chronic short-sleeping flies, such as the array of mutants that have been isolated in *Drosophila*^36–40^, may serve as a less stress-confounded model system for studying the relationship between sleep defects and oxidative stress. In addition to our lab, other groups have examined genetically short-sleeping fruit flies in the context of oxidative stress. One group observed that mitochondrial ROS serve as homeostatic regulatory elements in sleep-promoting neurons^41^, while another found an accumulation of ROS in the gut of flies and mice lacking sleep^42^. Despite these important findings, the core question remains: does lack of sleep induce pathology in a variety of tissues because of a specific mechanistic role in oxidative stress homeostasis? Ultimately, it is crucial to understand how sleep functions to promote health in a society that has become more chronically short-sleeping due to shift work, sleep disorders, and 24-hour access to dietary stimulants, fluorescent lights, and brightly-lit electronic screens.^43–46^

Here we utilized a diverse group of chronically short-sleeping *Drosophila* mutants and employed hyperoxia as an induced, acute oxidative stress to characterize the physiological state of short-sleeping mutants during oxidative stress. Using RNA-sequencing, we found that chronic short-sleeping mutants exhibit a typical transcriptional oxidative stress response relative to controls. Furthermore, we identified changes in gene expression that suggest a shift in metabolism from glycolysis towards lipid metabolism under hyperoxia. To test if short-sleeping mutants are more adversely affected by this hyperoxia-induced metabolic shift, we found that short-sleeping mutants are more sensitive to starvation than controls. Metabolomics revealed that hyperoxia led to reduced levels of long-chain fatty acids and lysophospholipids in short-sleeping mutants relative to controls, suggesting a defect in lipid metabolism. We found no specific defects in lipid metabolism common to short-sleeping mutants, such as triacylglyceride storage, respiration rate, or change in cuticular hydrocarbon profile (major products of fatty acid metabolism, similar to human sebum). Cuticular hydrocarbons are critical for protection against dehydration; we did find that all short-sleeping mutants tested are sensitive to dehydration, suggesting a common and general defect in fatty acid metabolism. Unexpectedly, we also found that, while flies with diet-induced obesity had no sleep or hyperoxia phenotypes, a genetic mutant (*brummer*) with obesity had a long-sleeping phenotype and was sensitive to hyperoxia. Because Brummer protein is a lipase that mobilizes fat stores, this suggests that genetic dysregulation of lipid catabolism could cause sensitivity to oxidative stress. Taken together, our work shows that chronic short sleep has a complex physiological impact on lipid metabolism, highlighting the possibility that metabolic defects may drive several common phenotypes of short sleeping mutants, including sensitivity to starvation, dehydration, and oxidative stress. These results are consistent with known associations between sleep disorders, metabolic defects, and diseases associated with oxidative stress, such as aging and neurological diseases.

## RESULTS

### Short-sleeping flies die more quickly than controls and exhibit increased nighttime sleep in hyperoxia

Previous methods for inducing acute exogenous oxidative stress (injection or feeding of paraquat or hydrogen peroxide)^47^ are either labor-intensive or dependent on feeding rate. Here, we tested if short-sleeping mutants are also sensitive to oxidative stress induced by hyperoxia, or 100% oxygen, which can be administered to many flies simultaneously with consistent dosage. Using *Drosophila* Activity Monitors (DAMs) in air-tight chambers to monitor for permanent inactivity as a proxy for death (**Fig. 1A-B**), we measured the survival of diverse short-sleeping flies in hyperoxia. We found that all short-sleeping mutants tested (*Hyperkinetic^Y^*, *Hk^Y^*; *insomniac*, *inc^1^* and *inc^2^*; *fumin*, *fmn*; *redeye*, *rye*)^37–39,48^ died significantly faster in hyperoxia than their genetic controls (**Fig. 1C**). Neither mutants nor controls died in normoxia (normal oxygen) conditions, in which compressed air is similarly administered in air-tight chambers. These results are consistent with our previous work showing significantly shorter survival of hyperoxia for short-sleeping *Cullin3* (*Cul3*) and *inc* (**Fig. S1A**) neuronal knockdown flies.^49^ Because these mutations and knockdowns affect sleep by different mechanisms, these results support the hypothesis that chronic sleep loss confers general sensitivity to an induced oxidative stress.

**Figure 1:**
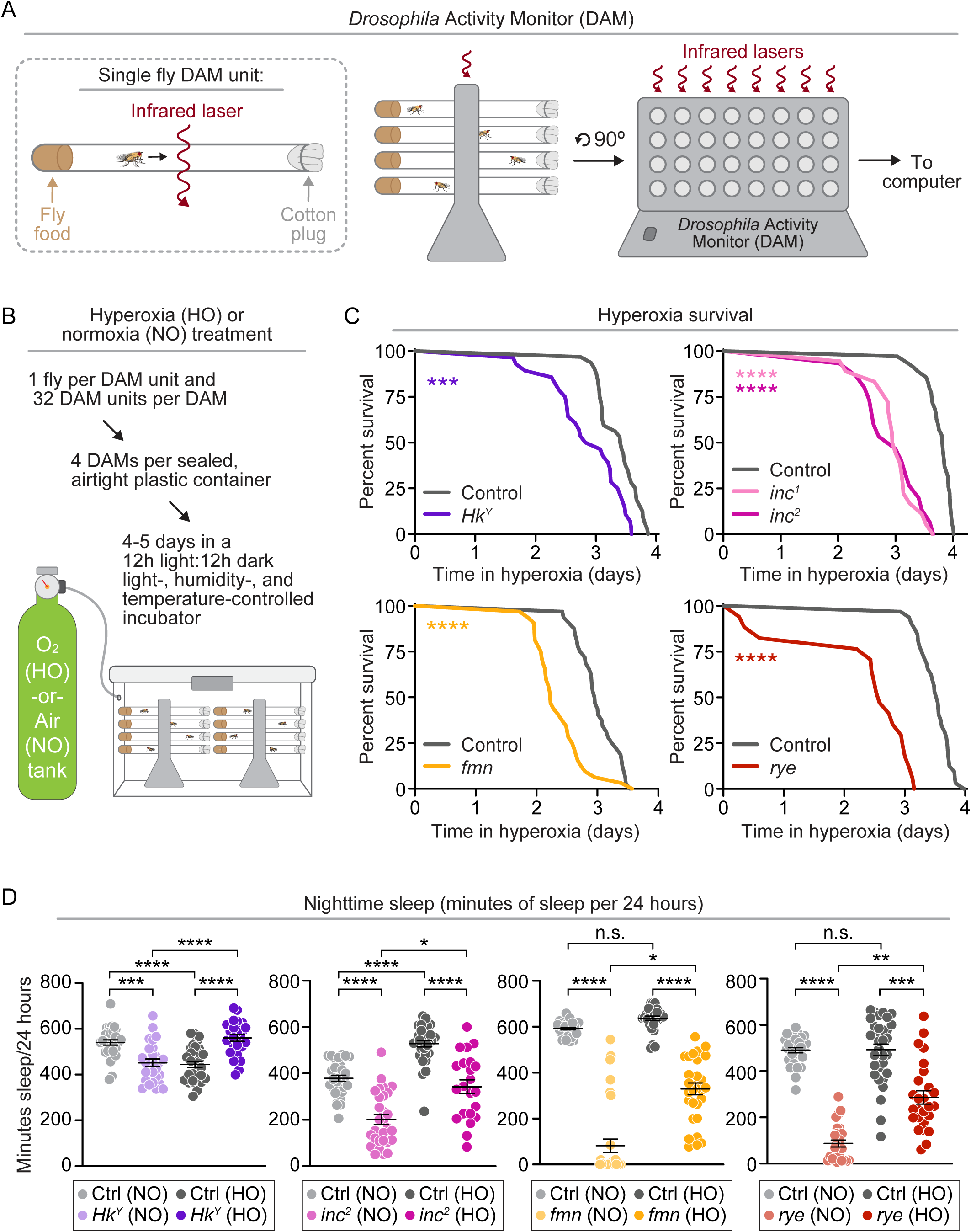
Short-sleeping mutants are sensitive to hyperoxia and have altered sleep. (A) Schematic of *Drosophila* Activity Monitors (DAMs) for measuring survival and sleep. (B) Schematic of DAMs in airtight plastic containers for normoxia or hyperoxia treatment. (C) Short-sleeping mutants *Hk^Y^* (p = 0.0005), *inc^1^* (p < 0.0001), *inc^2^* (p < 0.0001), *fmn* (p < 0.0001), and *rye* (p < 0.0001) die significantly faster than their respective controls in hyperoxia. (D) Short-sleeping mutants *Hk^Y^* (p = 0.0005), *inc^2^* (p = 0.0159), *fmn* (p = 0.0391), and *rye* (p = 0.0048) exhibit a significant increase in nighttime sleep in hyperoxia, as does the control for *inc^2^* (p < 0.0001), while *Hk^Y^*’s control loses sleep in hyperoxia (p < 0.0001). P-values were obtained by Log-rank test (C), Brown-Forsythe and Welch ANOVA test with Dunnett’s T3 multiple comparisons test (D), and Kruskal-Wallis test with Dunn’s multiple comparisons test (D). Averages are shown with error bars representing SEM.

We hypothesized that diverse short-sleeping mutants might have a common physiological defect due to chronic short sleep that drives their common sensitivity to oxidative stress. We tested if the sleep architecture of short-sleeping mutants exhibited a common alteration due to hyperoxia. Though both our *insomniac* alleles, *inc^1^* and *inc^2^*, are sensitive to oxidative stress, we focused on *inc^2^* mutants as *inc^2^* is an insertion mutation that affects only the *insomniac* gene, while *inc^1^* contains a deletion affecting both *insomniac* and a neighboring gene. We subjected four short-sleeping mutants and their genetic controls in DAMs to constant hyperoxia and analyzed the sleep of these flies in 24-hour increments from 0 to 72 hours of treatment using PySolo^50^, removing any flies that died during this time from the sleep analysis. This analysis revealed few common patterns in the sleep parameters of controls or short-sleeping mutants in hyperoxia (**Fig. S1**). The only common pattern exhibited was that all four short-sleeping mutants had significantly increased nighttime sleep relative to their genetic controls in hyperoxia (**Fig. 1D**).

### Short-sleeping flies can induce a normal transcriptional oxidative stress response

We hypothesized that the sensitivity to oxidative stress observed for short-sleeping mutants could stem from a common defect in inducing an oxidative stress response. The major oxidative stress response pathway (the Nrf2/Keap1 pathway) is transcriptionally regulated^51^; in oxidative stress conditions Nrf2 binds antioxidant response elements upstream of oxidative stress response genes. To test if short-sleeping mutants are defective in activating this pathway, we performed RNA-sequencing (RNA-seq) analysis on two short-sleeping mutants (*Hk^Y^* and *inc^2^*) and their genetic controls in both hyperoxia and normoxia conditions at four timepoints (6h, 12h, 18h, and 24h into hyperoxia or normoxia treatment) (**Fig. 2A**). Contrary to our hypothesis, we found that these short-sleeping mutants had similar or higher expression of Nrf2/Keap1 pathway genes relative to their controls (**Fig. S2A**). This result suggests that short-sleeping flies can induce an appropriate transcriptional antioxidant response; that is, a deficit in Nrf2/Keap1 activated transcription does not appear to be responsible for their increased sensitivity to oxidative stress. While short-sleeping mutants appeared to induce a normal transcriptional oxidative stress response, we wanted to test this response at the functional level. Using metabolomics (see Materials & Methods), we measured both reduced glutathione (GSH) and oxidized glutathione (GSSG) in several short-sleeping mutants and controls after 24 hours of normoxia or hyperoxia treatment. The ratio of reduced to oxidized glutathione (GSH:GSSG) can be used to measure oxidative stress levels^52^, with a lower GSH:GSSG ratio indicating more GSSG and therefore increased cellular oxidative stress. We found that, both at baseline and in hyperoxia, while *inc^2^* and *fmn* had significantly lower GSH:GSSG ratios relative to their controls, *Hk^Y^* had significantly higher or similar GSH:GSSG ratios relative to its control (**Fig. S2B**). These results suggest that *inc^2^* and *fmn* may experience higher levels of oxidative stress or *Hk^Y^* may be defective in its response to oxidative stress, but these defects are not common across these three short-sleeping mutants. Thus, chronic short sleep does not appear to cause sensitivity to oxidative stress because of a common failure to mount an oxidative stress response.

**Figure 2:**
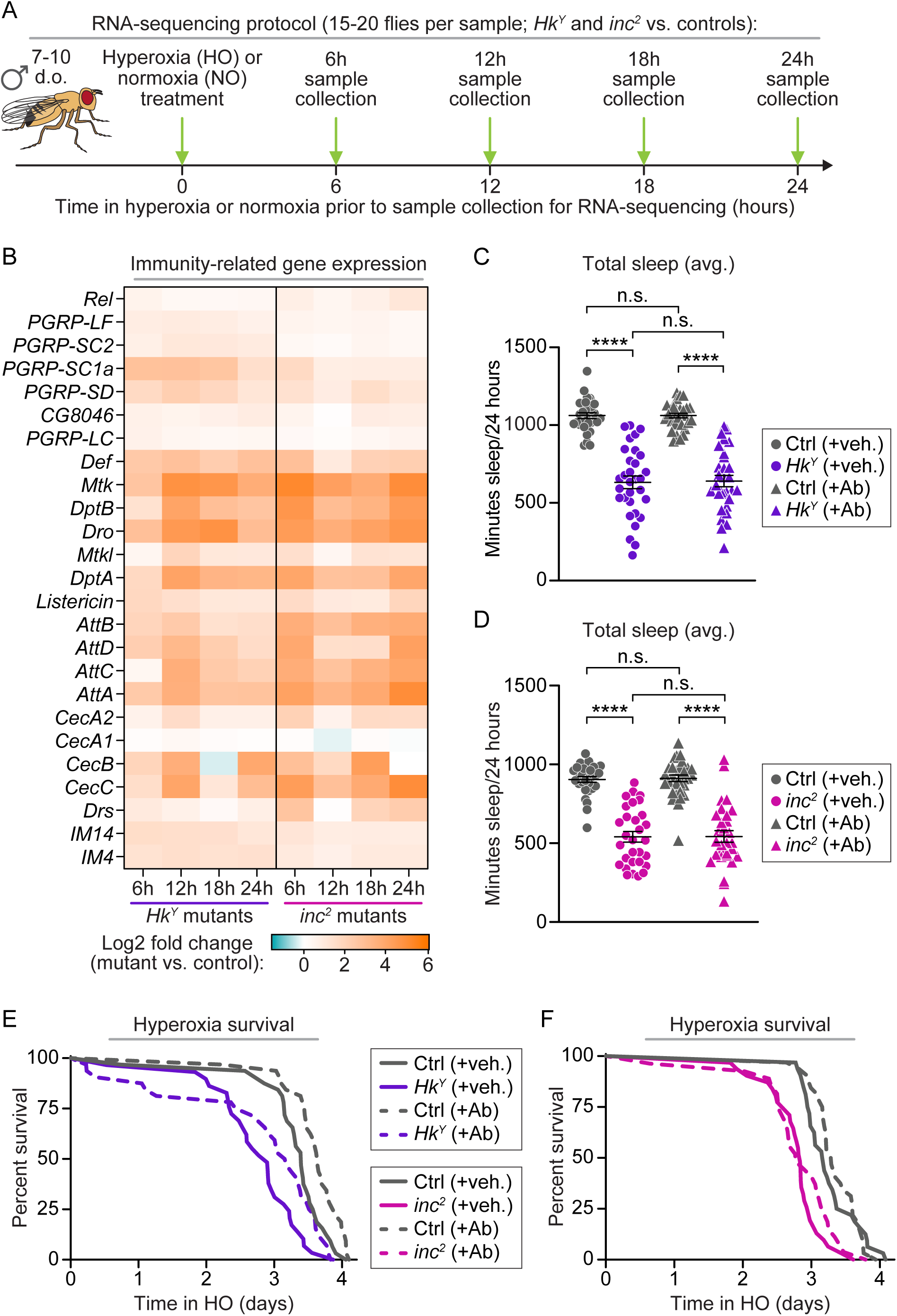
RNA-sequencing revealed higher expression of immune genes in short-sleeping mutants but sleep and oxidative stress sensitivity phenotypes are not changed by antibiotic feeding. (A) Schematic depicting the RNA-sequencing experimental workflow to assess transcription in young adult *inc^2^* and *Hk^Y^* flies in normoxia or hyperoxia compared to their respective controls. (B) Heatmap depicting the Log2 fold change of short-sleeping mutant differential gene expression for select immunity-related genes in normoxia compared to its genetic control across four timepoints. (C) Short-sleeping mutants *Hk^Y^* (p < 0.0001) and (D) *inc^2^* (p < 0.0001) are still significantly short-sleeping after being fed antibiotics from eclosion for 7 to 10 days. (E) Additionally, *Hk^Y^* (p < 0.0001) and (F) *inc^2^*(p = 0.0005) are still significantly more sensitive to hyperoxia relative to controls after antibiotic feeding. P-values were obtained by Brown-Forsythe and Welch ANOVA test with Dunnett’s T3 multiple comparisons test (C), Kruskal-Wallis test with Dunn’s multiple comparisons test (D), and Log-rank test (E-F). Averages are shown with error bars representing SEM.

### Short-sleeping flies exhibit chronic inflammation related to their gut microbiome, which is not responsible for their sensitivity to exogenous oxidative stress

A major category of differentially expressed genes identified in our RNA-seq analysis were immune response genes, particularly antimicrobial peptides (AMPs), which were upregulated in short-sleeping mutants relative to controls at baseline (**Fig. 2B**). We set out to determine whether this transcriptional inflammatory signature of the short-sleeping mutants was responsible for their increased sensitivity to oxidative stress. Low-level chronic immune stimulation is associated with microbiota load and oxidative stress, primarily in the fly gut^53^, and is inhibited by antibiotic treatment. We examined oxidative stress levels in the guts of short-sleeping mutants and their controls with and without antibiotic treatment. After 24 hours of normoxia or hyperoxia treatment, we dissected guts and stained them with dihydroethidium (DHE), an indicator of superoxide anion which is a main reactive oxygen species that contributes to oxidative stress. In normoxia, we found that DHE levels were higher in the guts of short-sleeping mutants relative to isogenic controls (**Fig. S3A-B**). Interestingly, hyperoxia increased the DHE levels of controls to those of short-sleeping mutants (**Fig. S3A-B**). As expected, antibiotic feeding significantly decreased the DHE levels of short-sleeping mutants to the same level as controls, both in normoxia and hyperoxia (**Fig. S3A-B**). These results are consistent with previous work showing that oxidative stress levels in fly guts can be associated with high bacterial load, a known factor for promoting ROS in the gut^54^, and show that the ROS levels of both sleep mutants and controls were reduced by antibiotic treatment.

Because antibiotic treatment reduced oxidative stress in the guts of short-sleeping flies, we next tested if these mutants were still short-sleeping and sensitive to hyperoxia without gut microbiome-related inflammation and increased ROS levels. We found that the short-sleeping mutants *Hk^Y^, inc^2^*, and their respective controls did not change their sleep behavior when fed antibiotics (**Fig. 2C-D**), suggesting that their inflammatory signature did not cause their short-sleeping phenotype. We also found that the antibiotic-treated short-sleeping mutants still exhibited significantly shorter survival in hyperoxia relative to their antibiotic-treated controls (**Fig. 2E-F**). As a note, antibiotic treatment extended hyperoxia survival for both *Hk^Y^* and its control (**Fig. 2E**), but not for *inc^2^* and its control (**Fig. 2F**). Taken together, these results suggest that gut microbiome-related inflammation and associated ROS levels do not mediate the increased sensitivity of short-sleeping mutants to acute, exogenous oxidative stress.

### Short-sleeping mutants exhibit dysregulated metabolism that may be impacted by the metabolic shift induced by oxidative stress

Metabolic genes constituted another major category of differentially expressed genes identified by our RNA-sequencing analysis. In hyperoxia relative to normoxia, for both short-sleeping mutants and controls and at all time points, we observed wide-spread downregulated expression of glycolytic genes and upregulated expression of fatty acid metabolic genes (**Fig. 3A-B**). This result suggested that, in the whole organism, hyperoxia causes a shift in metabolism away from carbohydrate metabolism and toward fat catabolism, a hypothesis supported by previous studies, mainly in tissue culture cells.^55–58^

**Figure 3:**
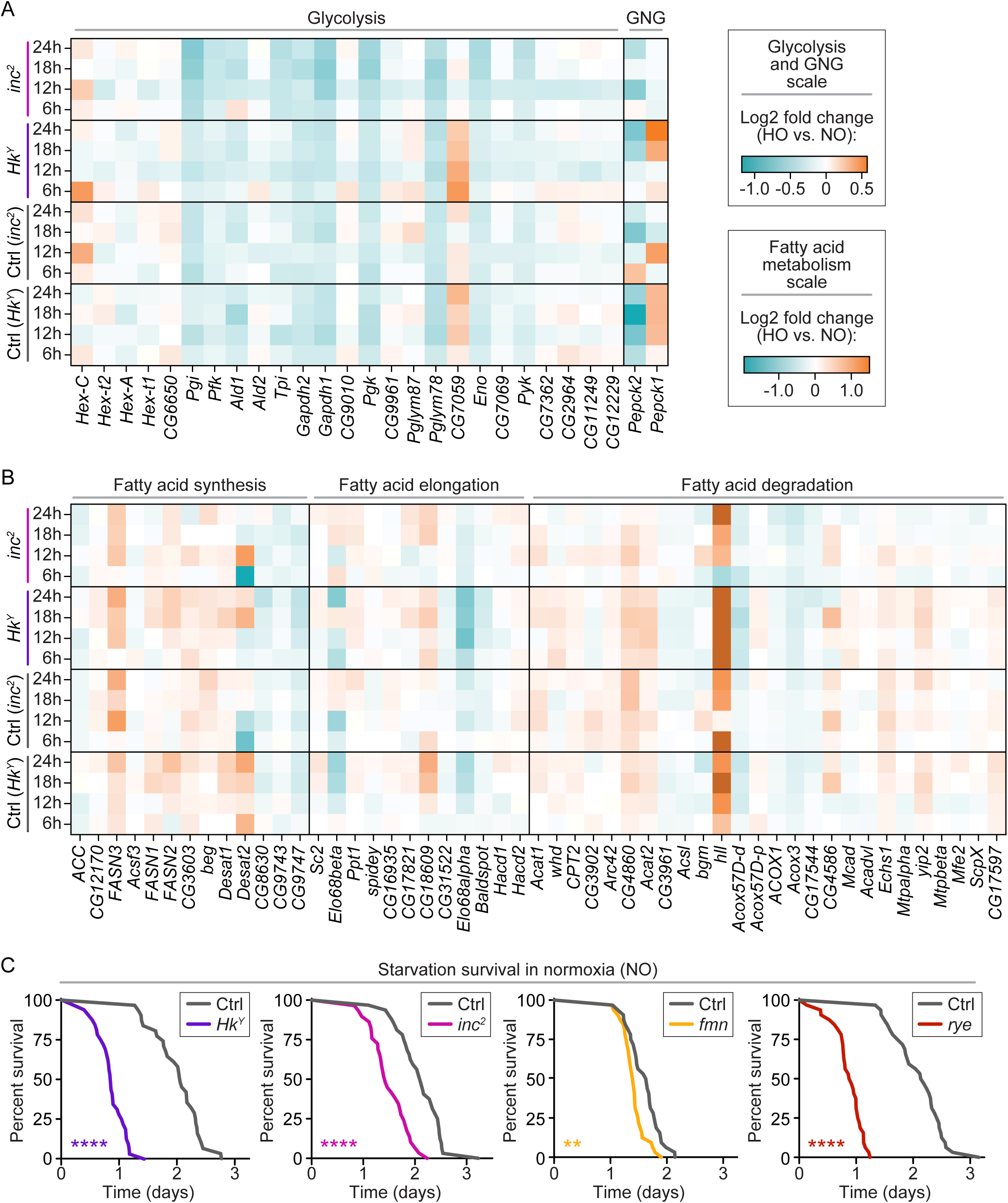
RNA-sequencing revealed differential expression of metabolic genes in hyperoxia, and between mutant and control at baseline, prompting investigation of short-sleeping fly metabolism. Heatmaps depicting the Log2 fold change of each genotype’s gene expression for select metabolic genes involved in (A) glycolysis and gluconeogenesis (GNG) and (B) fatty acid metabolism at four separate timepoints in hyperoxia compared to normoxia conditions. (C) All short-sleeping mutants, *Hk^Y^* (p < 0.0001), *inc^2^* (p < 0.0001), *fmn* (p = 0.0020), and *rye* (p < 0.0001) starve to death significantly faster than their respective controls. For metrics used the log2fold change, refer to methods. P-values were obtained by Log-rank test (C).

Sleep and fasting are two other biological states known to shift metabolism away from glycolysis and toward fat catabolism.^52,59–61^ We hypothesized that short-sleeping mutants may have difficulty dealing with the consequences of this metabolic shift. To test this, we measured survival time in starvation conditions and found that short-sleeping mutants died significantly faster than isogenic controls (**Fig. 3C**). This result suggests that short-sleeping mutants have a common metabolic defect and further suggests that their sensitivity to oxidative stress could reflect a broader defect in fasting metabolism.

In the simplest hypothesis, oxidative stress might itself induce starvation for both short-sleeping mutants and controls, leading to the observed shift in metabolic gene expression and shorter survival of short-sleeping mutants. To test this possibility, we first measured the food consumption of short-sleeping mutants and controls in both normoxia or hyperoxia conditions. In normoxia, we found that *Hk^Y^*, *fmn*, and *rye* consumed significantly more food compared to their respective genetic controls, while *inc^2^* ate the same amount as its control (**Fig. S4A**). In hyperoxia, we found that all eight genotypes (four short-sleeping mutants and their isogenic controls) ate less than they did in normoxia (**Fig. S4A**). Thus, while short sleep did not have a common impact on baseline feeding behavior, hyperoxia did have a common impact on all genotypes.

To test if flies in hyperoxia are simply starving to death, we tested the effect of combined hyperoxia and starvation conditions on survival time. We found that the four control genotypes died significantly faster with the combined stress of starvation and hyperoxia, compared to starvation alone or hyperoxia alone (**Fig. S4B**). That is, for control flies, these stresses are clearly additive in their reduction of survival time. The results were more variable for the short-sleeping mutants. For *inc^2^*, there was no change in survival when hyperoxia and starvation were combined compared to starvation alone, while *fmn* and *rye* died at a faster rate under the combined stresses relative to either stress alone. Unexpectedly, for *Hk^Y^*, the combined stress of hyperoxia and starvation increased survival time relative to starvation alone. The lack of a consistent additive phenotype for short-sleeping mutants contrasted with the consistent additive phenotype of control flies under combined oxidative stress and starvation. Taken together, these results suggest that, while hyperoxia may induce mild fasting, hyperoxia does not induce starvation and, while short-sleeping mutants have common sensitivity to starvation, this phenotype has complex effects when combined with hyperoxia.

### Global metabolomics of short-sleeping flies in hyperoxia reveals a significant decrease in lysophospholipids and fatty acids

To further investigate the metabolic defects of short-sleeping mutants relative to controls, we performed global metabolomics on *Hk^Y^*, *inc*^2^, *fmn* and their respective genetic controls in conditions of normoxia or hyperoxia for 24 hours. We hypothesized that if hyperoxia (and the accompanying shift in metabolism) exacerbates a defect in metabolic homeostasis common to the short-sleeping mutants, we would see a change in metabolites during hyperoxia common to the short-sleeping mutants relative to their controls. Strikingly, there were only two groups of metabolites that changed specifically during hyperoxia for all three short-sleeping mutants relative to controls: lower levels of long-chain fatty acids (**Fig. 4A**) and lysophospholipids (**Fig. 4B**). These results suggest that specific aspects of fat catabolism, particularly those producing fatty acids and lysophospholipids, are dysregulated in short-sleeping mutants during acute oxidative stress.

**Figure 4:**
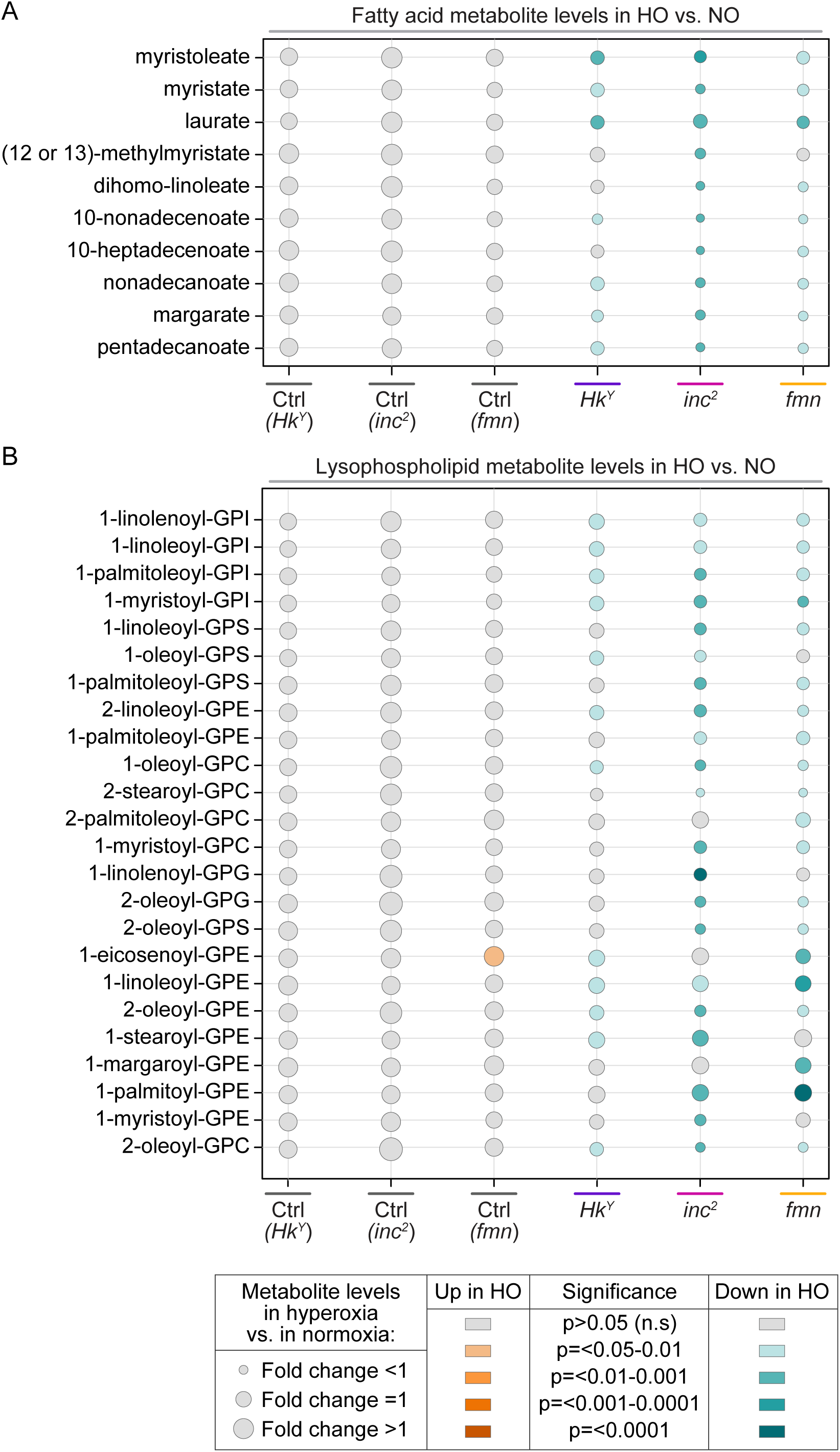
Metabolomics highlighted significant metabolite changes to short-sleeping flies in hyperoxia. (A-B) Bubble plots depicting the fold change, direction (up-ordown-regulated) and statistical significance of metabolite differences for each genotype in hyperoxia compared to normoxia. The plots reveal significantly downregulated fatty acids (A) and lysophospholipids (B) in all short-sleeping mutants in hyperoxia, but no difference for controls in hyperoxia. P-values were obtained by t test.

### Short-sleeping mutants did not exhibit a common defect in lipid metabolism in hyperoxia

To investigate if short-sleeping mutants have other common defects in lipid metabolism, we characterized several metabolic parameters, including their basal fat stores, fat loss over time in hyperoxia, respiration rates, and cuticular hydrocarbon levels. First, we hypothesized that short-sleeping mutants might be sensitive to starvation because they have lower fat stores or are impaired in fat utilization relative to controls. Triacylglycerides (TAGs) are the main fat storage molecule in both humans and flies and can be quantitatively measured using thin-layer chromatography (**Fig. 5A**). To test for a common defect, we measured TAG levels in four short-sleeping mutants and their genetic controls in both hyperoxia and normoxia conditions. While two out of four mutants (*Hk*^Y^ and *inc^2^*) did have significantly lower TAG stores than their controls in both conditions (**Fig. 5B, S5A**), one mutant (*rye*) was not significantly different from its control in either condition (**Fig. 5B**) and one mutant (*fmn*) had significantly increased TAG stores in both conditions (**Fig. S5A**). Moreover, all four mutants lost TAG levels during hyperoxia similarly with their controls. That is, we did not observe common differences in TAG levels between short-sleeping mutants and controls, suggesting that the sensitivity of short-sleeping mutants to oxidative stress is not due to a common defect in TAG storage or mobilization.

**Figure 5:**
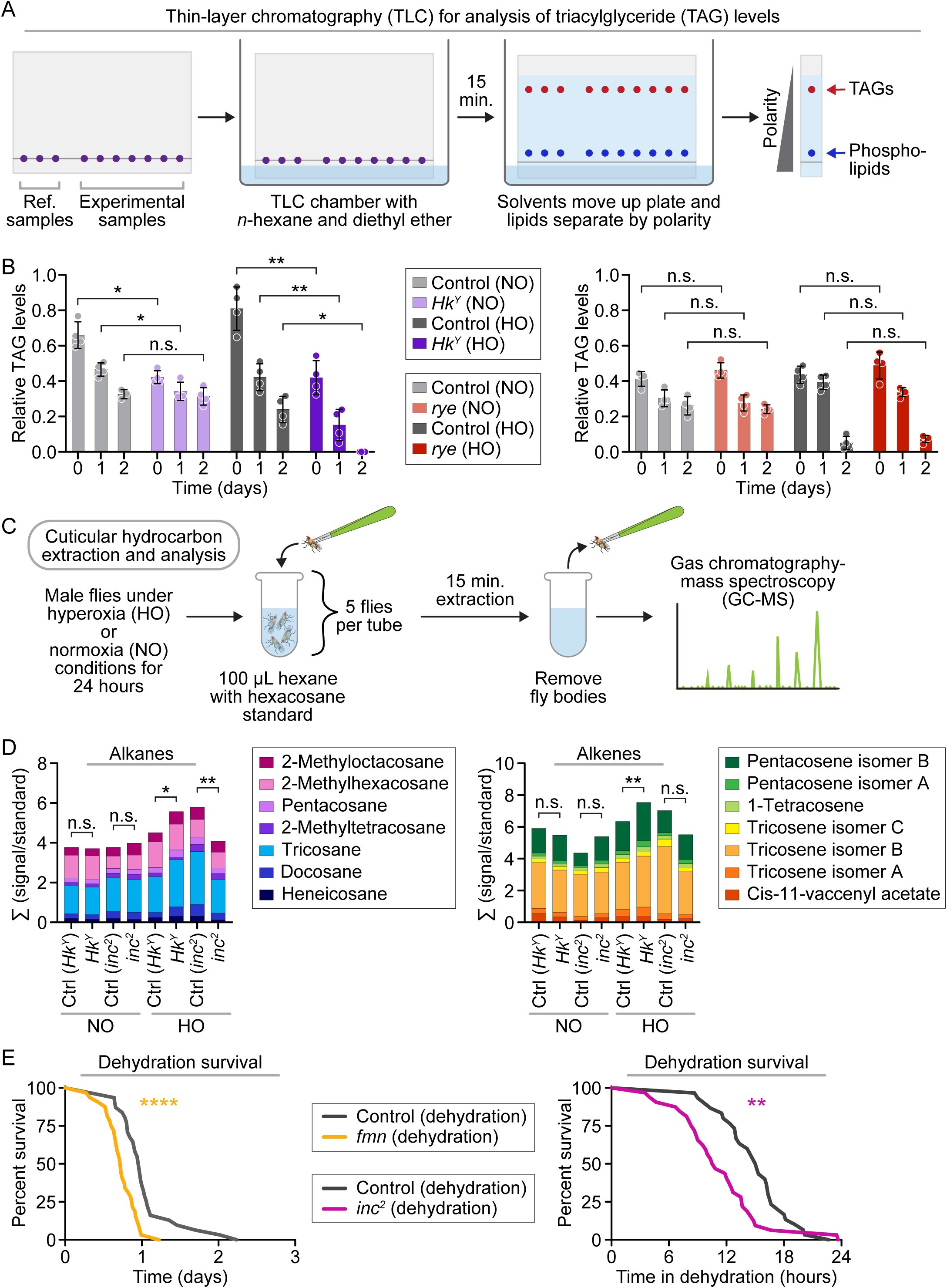
Short-sleeping mutants exhibit variable triacylglyceride stores, kinetics of usage, and trend in Cuticular Hydrocarbon compound profile. (A) Schematic depicting the TLC (Thin-layer chromatography) set up for testing Triacylglyceride(TAG) levels. (B) TLC analysis revealed that *Hk^Y^* has more fat (control NO 0 vs *Hk^Y^* NO 0, p= 0.0124) and loses TAGs at a faster rate than its control in hyperoxia (control NO 24 vs *Hk^Y^* NO 24, p= 0.9593, control HO 48 vs *Hk^Y^*HO 48, p= 0.0180) but *rye* and its control has similar basal level of TAGs (control NO 0 vs *rye* NO 0, p= 0.6173) and loses fat in similar level control in hyperoxia (control NO 24 vs *rye* NO 24, p= 0.7989, control HO 24 vs *rye* HO 24, p= 0.1995). Dots represent biological replicates of 10 male flies. (C) Schematic depicting the Cuticular Hydrocarbon extraction and analysis experimental workflow. (D) GC-MS analysis of sleep mutant cuticular hydrocarbon profiles showed that there is overall increase in alkane in HkY after 24 hours of hyperoxia. No common significant trend was observed in change in overall or individual alkanes or alkene compound was found. (E) Short sleeping mutants *fmn* and *inc^2^* showed sensitivity to dehydration. P-values were obtained by Brown-Forsythe and Welch ANOVA test with Dunnett’s T3 multiple comparisons test (B), Wilcoxon test (D), or Log-rank test (E). Averages are shown with error bars representing SEM.

Second, we examined respiration rates. Overall metabolic rate is known to be higher during wake than sleep^62^ and short-sleeping mutants are awake longer than their controls. Thus, we hypothesized that short-sleeping mutants might be sensitive to starvation and oxidative stress because of higher metabolic rates at baseline or an inability to maintain or increase metabolic rate under hyperoxia. To test this, we measured CO_2_ production as a proxy for metabolic rate using soda lime-based respirometers (see Materials & Methods) (**Fig. S5B)**. Measurement of CO_2_ production for all genotypes at ZT 1-3 after 24 hours of normoxia or hyperoxia treatment identified no common phenotypes across genotypes (**Fig. S5C**): only *rye* had significantly higher respiration than its control at baseline and in hyperoxia, while *Hk^Y^* was the only mutant to exhibit a significant decrease in its respiration rate in hyperoxia. These results suggest that chronic short sleep does not cause a common change in the overall metabolic rate, either at baseline or in hyperoxia, that drives sensitivity to starvation or oxidative stress.

Third, we performed a quantitative analysis of cuticular hydrocarbon (CHC) profiles. Deposited on the external cuticular surface, CHCs are one of the main end products of fatty acid metabolism in *Drosophila* and have two major functions: mediating chemical communication between individuals (*i.e.*, as cues for courtship and aggression) and, like human sebum, forming a waxy barrier that protects against loss of moisture or desiccation. We hypothesized that if short-sleeping mutants had altered lipid metabolism, they would exhibit altered CHC composition and/or altered resistance to dessication. To measure CHC composition, we extracted lipid content specifically from the external cuticular surface of two short-sleeping mutants and controls after 24 hours of normoxia or hyperoxia and analyzed by gas chromatography mass spectroscopy (GC-MS) **(Fig. 5C)**. Overall, hyperoxia increased total alkene and alkane CHCs for both short-sleeping mutants and controls relative to normoxia; this result is consistent with increased fatty acid metabolism during hyperoxia. However, there were very few consistent differences in individual CHC species between *Hk^Y^* or *inc^2^* mutants and their controls **(Fig. 5D).** Only two CHC species (pentacosene isomers A and B) were increased in hyperoxia for both *Hk^Y^* and *inc^2^* mutants relative to their controls. These results suggest that chronic short sleep does not cause a common defect in CHC composition or levels.

Finally, to test broad CHC functional differences between short-sleeping mutants and controls, we tested their dehydration resistance. Previous studies found that variations in CHC components across flies directly affect their desiccation resistance.^63^ We modified the protocol from Wang et al. to develop DAM-based experimental setups testing either dehydration survival (no food, no water) or desiccation survival (no food, no water, plus silica gel desiccant to actively remove moisture from the air). We found that all four short-sleeping mutants died significantly more quickly than their isogenic controls in either dehydration or dessication conditions (**Fig. 5E, Fig S5B)**, suggesting an overall defect in CHCs that was not apparent from examining individual cuticular hydrocarbon species. Together, these results suggest that a common aspect of lipid metabolism (independent of fat storage, respiration rates, and individual CHC species) is dysregulated in short-sleeping mutants and that this common defect in lipid metabolism might mediate their common sensitivities to oxidative stress, starvation, and dehydration.

### Genetic lipid metabolic defect (*brummer*), but not diet-induced obesity (high sugar diet), affected sleep and sensitivity to hyperoxia

To test if altering fat stores or inducing defects in lipid metabolism could alter sleep or sensitivity to hyperoxia, we tested two types of obese flies: those raised on high-sugar diet (0.7 M Sucrose for 10 days) and *brummer* mutants, which lack a conserved TAG lipase (adipose triglyceride lipase or ATGL in humans) that catalyzes the first step of lipolysis or fat mobilization. Consistent with previous reports, both high-sugar diet flies and *brummer* mutants had increased fat stores relative to controls, as measured by thin-layer chromatography (**Fig. 6A**). While high-sugar diet flies exhibited no changes in sleep, *brummer* mutants were long sleepers and exhibited increased total sleep time (**Fig. 6B**). In our previous publication, we had found that all short sleepers tested were sensitive, and all long sleepers resistant, to oxidative stress relative to controls. Consistent with this, high-sugar diet flies, which had normal sleep, exhibited no difference in survival time in hyperoxia relative to low-sugar diet controls (**Fig. 6C**). In contrast, long sleeping *brummer* mutants were not resistant to hyperoxia as expected, but sensitive to hyperoxia (**Fig. 6F**). These results suggested that merely increasing fat stores through high-sugar diet is not sufficient to alter sleep or sensitivity to hyperoxia but a genetic defect in lipid metabolism can impact both sleep homeostasis and sensitivity to hyperoxia.

**Figure 6:**
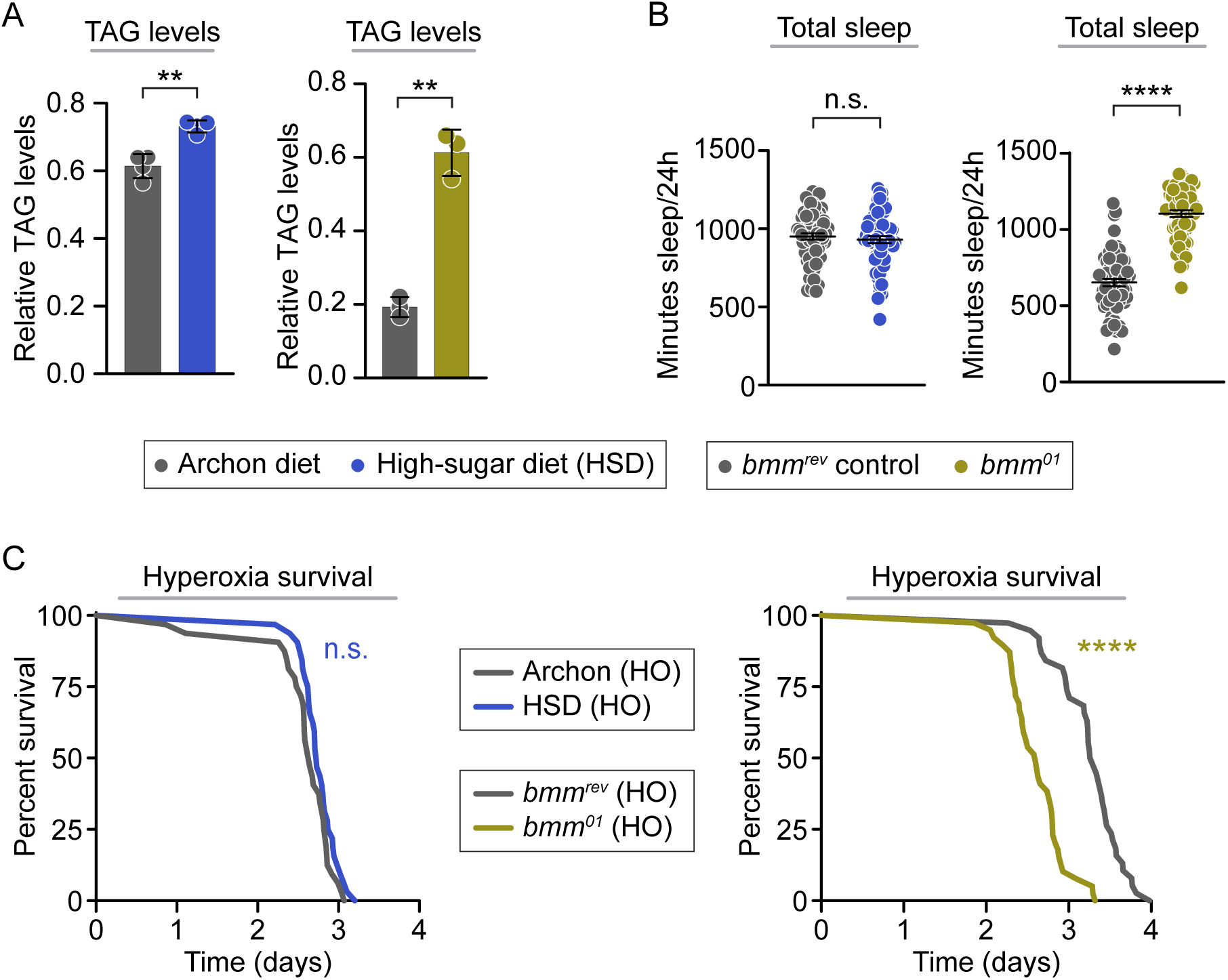
Various lipid mutants exhibit range of sleep and hyperoxia sensitivity. (A) Flies raised in high sugar diet post eclosion for 10 days that has higher fat level and *Bmm* mutants have significantly higher TAG level than its control, *bmmrev* (p < 0.0001). (B) High sugar diet flies have similar total sleep to control (p= 0.5279). *Bmm* mutants sleeps significantly more overall than its controls (p < 0.0001). (C) High sugar diet flies had no difference in terms of hyperoxia survival (p=0.1053). *Bmm* flies die faster in hyperoxia (p < 0.0001) compared to its controls. P-values were obtained by or one-way ANOVA test (A), Mann-Whitney Test (B) and Log-rank test (C). Averages are shown with error bars representing SEM.

## DISCUSSION

Based on our lab’s previous work establishing a bidirectional relationship between sleep and oxidative stress in fruit flies^47^, we set out to characterize the physiological state of short-sleeping mutants under oxidative stress. We first confirmed that, like their sensitivity to other types of oxidative stress, diverse short-sleeping mutants were significantly more sensitive to hyperoxia than isogenic controls. RNA-seq analysis revealed that two different short-sleeping mutants induced typical transcriptional oxidative stress responses, similar to those of isogenic control flies. Metabolomics supported this, showing that short-sleeping mutants did not have a common defect in glutathione ratios. Our RNA-seq analysis led to two additional findings: an inflammatory transcriptional signature in short-sleeping mutants at baseline and differential expression of metabolic genes for both short-sleeping mutants and controls in hyperoxia, suggesting a metabolic shift away from carbohydrate catabolism toward fat catabolism. Because antibiotic treatment did not impact sleep or oxidative stress sensitivity of short-sleeping mutants, we focused on the metabolic phenotype. Though our previous data suggested that neuronal *inc* RNAi knockdown did not cause sensitivity to starvation, here we found that all short-sleeping mutants tested (including neuronal *inc* RNAi knockdown flies) were more sensitive to starvation than isogenic controls. Using metabolomics, we found that three different short-sleeping mutants had common reductions in long-chain fatty acid and lysophospholipid species in hyperoxia conditions. Searching for specific defects in lipid metabolism, we compared TAG storage levels, cuticular hydrocarbons profiles, and respiration rates of short-sleeping mutants and isogenic controls in normoxia and hyperoxia. We found no defects in these aspects of lipid metabolism that were common across all three short-sleeping mutants, though we did find that short-sleeping mutants were more sensitive to dehydration than controls, suggesting an overall common defect in cuticular hydrocarbon function. To test if changes in lipid metabolism could alter sleep or sensitivity to oxidative stress, we increased fat stores either through high sugar diet or genetic mutation. We found that diet-induced obesity (flies on high sugar diet) had no effect, but genetic obesity (lipid mobilization mutant *brummer*) caused longer sleep and increased sensitivity to hyperoxia. This phenotype overturns the paradigm established by our previous work and suggests the intriguing possibility that lipid homeostasis is monitored by the sleep homeostat. That is, if sleep regulates lipid homeostasis, then the mechanism regulating sleep does not monitor total lipid storage but instead appears to monitor lipid mobilization kinetics. Taken together, the results here extend the previously published work on sleep and oxidative stress and underscore the need to understand the role of sleep in lipid metabolism, as this could drive the known pathological associations between sleep disorders, obesity, and diseases of oxidative stress.

Of particular interest is our finding that oxidative stress caused significant global reductions in multiple species of long chain fatty acids and lysophospholipids only in short-sleeping mutants. While we are not the first lab to perform metabolomics on genetically short-sleeping *Drosophila* mutants, we are the first to do so in oxidative stress conditions. Bedont *et al*. (2023)^64^ performed global metabolomics and lipidomics with Metabolon on short-sleeping mutant heads. They found that short-sleeping mutants accumulated polyamines and exhibited inefficient nitrogen excretion, leading them to characterize an overall nitrogen stress sensitivity in short-sleeping mutants. In our whole-body metabolomics with Metabolon, we did not observe clear differences in polyamine or nitrogen-related metabolism in our short-sleeping flies relative to their genetic controls. The differences in metabolite patterns in our two datasets may be due to the different tissues assayed—just heads (Bedont analysis) versus entire bodies (this analysis). It is known that metabolism can differ between different tissues.^65^ For example, Wilinski et al found differential gene expression and metabolite presence in heads versus bodies when flies were starved.^66^ It may be that short sleep has different metabolic consequences on different tissues of the fly. Tissue-specific metabolomics of short-sleeping flies versus their genetic controls in normoxia and hyperoxia could resolve differences across studies.

As hyperoxia was critical for identifying this defect in lipid-based metabolites, we examine here our use of hyperoxia. In hyperoxia, both short-sleeping mutant and control flies underwent a general shift in gene expression away from carbohydrate metabolism and toward fat catabolism. This hyperoxia-induced shift in metabolism is supported by previous studies. Hyperoxia treatment increased ROS and reduced glucose uptake in adipose tissue culture cells^55^ and decreased glycogenolysis in muscle during aerobic exercise by decreasing the system’s reliance on carbohydrate and promoting fat oxidation^56^. Metabolomic analysis of *Drosophila* after paraquat exposure, another method of inducing oxidative stress, also found alterations in metabolites related to carbohydrate metabolism and lipid catabolism.^58^ Conversely, hypoxia (low oxygen) is known to inhibit oxidative phosphorylation and fatty acid metabolism in tissue-culture cells.^67^ It is possible that the hyperoxia-induced shift in metabolism is not due to oxidative stress *per se* but simply due to the increase of oxygen flux through the election transport chain. Still, this assay proved useful as it allowed us to identify altered fatty acid metabolism as a common phenotype of short-sleeping mutants.

We have now identified several phenotypes related to fatty acid metabolism that are common to short-sleeping mutants: sensitivity to starvation, sensitivity to dehydration, and reduced levels of long-chain fatty acids and lysophospholipids in hyperoxia. These phenotypes demonstrate that sensitivity to oxidative stress is not the sole nor unique phenotype of short-sleeping mutants and suggest that short sleep has broad physiological impact, particularly for physiological functions dependent on lipid metabolism. It is possible that short sleep independently causes multiple phenotypes; alternatively, short sleep might cause a major physiological defect that drives these phenotypes or one of these phenotypes might drive the other phenotypes.

It should be noted that, while we identified defects in specific lipid species and physiological functions that were common across diverse short-sleeping mutants, we did not identify a defective step of lipid metabolism that was common across diverse short-sleeping mutants. Our approach has rested on two fundamental assumptions: first, that one function of sleep is to regulate a specific cellular or physiological mechanism; and second, if so, that all short-sleeping mutants would have common defects in that specific cellular or physiological mechanism. Based on these results, we note the possibility that sleep may have broad and diverse effects on physiology and that diverse short-sleeping mutants might have common sensitivities to stress due to diverse mechanistic defects. In other words, different short-sleeping mutants might be sensitive to oxidative stress for different reasons or have reduced lysophospholipids during oxidative stress because of defects at different stages of lipid metabolism. To paraphrase Dr. Kyunghee Koh (who paraphrases Dostoevsky): Perhaps all healthy sleepers are alike, but every unhealthy sleeper is unhealthy in its own way. Given the crucial importance of sleep for human health, future research is needed to distinguish between these possibilities and determine the specific physiological functions of sleep.

## MATERIALS & METHODS

### Fly strains and rearing conditions

| Genotype | Source |
| --- | --- |
| $w^{1118}; w; w$ | Daniel Bushey |
| $w^{1118}; Hk^Y; w; w$ | Daniel Bushey |
| $inc^1, w^+; w; w$ | Nick Stavropoulos |
| $inc^2, w^{1118}; w; w$ | Nick Stavropoulos |
| $w^{1118}; w; w$ | Nick Stavropoulos |
| $w^{1118}; w; w$ | Rob Jackson |
| $w^{1118}; fmn/CyO; w$ | Rob Jackson |
| $w; w; w/TM6cSb$ | Amita Sehgal |
| $w; w; rye/TM6cSb$ | Amita Sehgal |
| $elavC155-gal4; w; w$ | Nick Stavropoulos |
| $w^-; UAS-inc.RNAi.1; w$ | Nick Stavropoulos |
| $w^-; w; w$ | Nick Stavropoulos |
| $w^{1118}; bmm^1/sb$ | Liz Rideout |
| $w^{1118}; bmm^{rev}$ | Liz Rideout |

All flies were raised on glucose food (Archon) in a temperature-(25°C) and humidity-(65%) controlled incubator with a 12-hour light-dark cycle. 7-to 10-day-old males were used for all experiments, unless otherwise noted.

### Sleep and activity analysis

7-to 10-day-old male flies entrained on a 12:12 light-dark cycle were placed in individual 5 mm plastic tubes containing food. Tubes were placed in TriKinetics *Drosophila* Activity Monitors (DAMs) to record their locomotor activity for 2 days in 12:12 light-dark. Beam break data were grouped into 1-min bins using DAM File Scan, and pySolo (Python-based software) was used to analyze sleep architecture and waking activity. Sleep was defined as a period of at least 5 min of inactivity.

### Normoxia and hyperoxia treatment

7-to 10-day-old male flies entrained on a 12:12 Light-Dark cycle were placed in vials or *Drosophila* Activity Monitors containing food, then placed inside of airtight chambers (Kent Scientific VetFlo™ Low Cost Induction Chamber) filled with medical air or 100% oxygen.

### Hyperoxia assay

7-to 10-day-old male flies entrained on a 12:12 Light-Dark cycle were placed in individual 5 mm plastic tubes containing food and loaded into *Drosophila* Activity Monitors. The monitors were placed inside of airtight chambers (Kent Scientific VetFlo™ Low Cost Induction Chamber), filled with 100% oxygen starting at ZT0 (“lights on”) until all flies were dead. Time of death was determined by a permanent loss of movement.

### RNA extraction and sequencing

Fifteen to twenty 7-to 10-day-old male flies were placed in each vial and treated with normoxia or hyperoxia. Vials were removed at 6, 12, 18, and 24 hours of treatment and flipped into Eppendorf tubes. RNA was extracted from samples by Columbia University Irving Medical Center’s Molecular Pathology Shared Resource. RNA samples were sequenced by JP Sulzberger Columbia Genome Center using a standard RNA-seq pipeline. Standard poly-A pull-down was used to enrich mRNAs from total RNA samples and RIBOZERO was used for ribosomal depletion to remove rRNAs from total RNA samples, followed by library construction using Illumina TruSeq chemistry. Clontech Ultra Low v4 kit was used for cDNA synthesis followed by NexteraXT.

Libraries were then sequenced using Illumina NovaSeq 6000, with multiplexed samples in each lane, which yielded a targeted number of paired-end 100 bp reads for each sample. RTA (Illumina) was used for base calling and bcl2fastq2 (version 2.19) for converting BCL to fastq format, coupled with adaptor trimming. A pseudoalignment was performed to a kallisto index created from transcriptomes (Human: GRCh38; Mouse: GRCm38) using kallisto (0.44.0). Differentially expressed genes under various conditions were tested using Sleuth or DESeq2, R packages designed to test differential expression between two experimental groups from RNA-seq counts data.

### Antibiotic feeding

Flies collected soon after eclosion were placed on vehicle-or antibiotic-supplemented food. The antibiotic cocktail was made of ampicillin (5 g), tetracycline (0.5 g), and rifamycin (2 g) in 100 mL of 50% ethanol to create a 100X concentration. Volumes were added to Archon glucose food to yield 1X concentration of antibiotic-supplemented food or equal volume of ethanol (vehicle)-supplemented food. Flies remained on their respective diet, flipped to fresh food every other day, through experimentation.

### DHE gut staining

Flies were anesthetized on ice and intestines were dissected in cold Schneider’s Medium (Thermo Fisher Scientific). Intestines were then incubated in 50 µM DHE (Invitrogen) in Schneider’s Medium for 5 min at room temp, then washed three times for 30 sec in Schneider’s Medium. DHE samples were quickly mounted in Schneider’s Medium. Intestines were imaged within 20 min on a Ziess LSM800 confocal microscope using identical exposure settings with 20x air objective (excitation 525 nm, emission 610 nm). Z-stacks spanning the entire posterior midgut were taken. Quantification of DHE was performed using FIJI^68^, in which mean fluorescent intensity values were recorded per Z-slice then averaged per gut. A minimum of 9 midguts were used for each genotype or condition.

### Starvation assay

Seven-to ten-day-old male flies were individually placed in 5 mm plastic tubes containing 1% agar and loaded into *Drosophila* Activity Monitors. Time of death was determined by complete loss of movement.

### Feeding assay

As previously described^69^, approximately 7-to 9-day-old flies were fed ^32^P-labeled Archon food for 24 hours. Scintillation counts of accumulated ^32^P were quantified for an average of 5 flies per sample to estimate the amount of food consumed.

### Triacylglyceride measurement using thin-layer chromatography

Each biological replicate sample was comprised of 10x 7-to 10-day-old male flies. Samples were homogenized in 125 µL of 2:1 chloroform:ethanol solution, then centrifuged for 15 min at 4°C. Lipids were then extracted and subsequently diluted to achieve concentrations within the linear range of our reference samples. Reference samples were prepared by pooling extracted lipids from 25 replicates of 20 females in 200 µL of 2:1 chloroform:ethanol, which are diluted to reach the linear range within a serial dilution series. 10 µL of each sample were loaded on a TLC plate (Sigma Aldrich Silica gel on TLC-PET foils 99577-25EA) and separated by polarity using a solvent solution (70 mL n-hexane, 30 mL diethyl ether, and 1 mL acetic acid). Plates were stained with 0.2% amido black (NAPHTHOL BLUE BLACK, Sigma Product N-3393-100G) in 1 M NaCl and imaged with an iBright™ FL1500 Imaging System. Images were analyzed using FIJI and quantified by measuring raw integrated density, subtracting individual lane background for each sample, and normalizing to the average of the reference samples.

### CO_2_ respiration assay

Measurement of CO_2_ respiration was performed as previously described.^70^ Ten 7- to 10-day-old male flies were placed in a sealed 1000 uL pipette tip-capillary respirometer (without anesthesia) containing 3 pellets of soda lime. Junctions of the respirometer were sealed tightly with Parafilm. Loaded respirometers were placed capillary side down in dH_2_O containing 0.1% blue food dye in a sealed glass chamber placed within a 25°C incubator. Several empty respirometers (containing no flies) were placed in the chamber, as well, to serve as a background control. Flies were tested for 1.5 to 2 hours at each circadian timepoint, and CO_2_ production rate was determined by the amount of liquid that had moved up the capillary per after, normalized to the background controls (empty respirometers). At least six groups of ten flies were used for each genotype/condition.

### Metabolomics

7- to 10-day-old male flies entrained on a 12:12 light-dark cycle were placed in vials for 24 hours of normoxia or hyperoxia treatment. After treatment, at ZT0 (“lights on”) flies were flipped into vials in groups of 70 males, weighed, and frozen at-80°C. Samples were shipped on dry ice to Metabolon for global metabolomics. Sample preparation, ultrahigh performance liquid chromatography-tandem mass spectroscopy, and bioinformatic analysis was performed by Metabolon. Metabolon utilized a a Waters ACQUITY ultra-performance liquid chromatography (UPLC) and a Thermo Scientific Q-Exactive high resolution/accurate mass spectrometer interfaced with a heated electrospray ionization (HESI-II) source and Orbitrap mass analyzer operated at 35,000 mass resolution. Welch’s two sample t-tests were used to determine significant alterations in metabolite levels between samples.

### Cuticular Hydrocarbon Analysis

Day 7 post-eclosion male flies entrained on a 12:12 light-dark cycle were placed in vials in groups of 10 flies per vial for 24 hours of normoxia or hyperoxia treatment. After the treatment, the flies were cold anesthetized and collected into an amber glass 2 mL vial (Agilent, P/N 5190-9590) fitted with a 150 µL glass insert (Agilent, P/N 5183-2088) and closed with a Teflon-lined septum cap (Agilent, P/N 5191-8160). Whole body liquid extraction was performed in a solution consisting of 90 µL of hexane and 10 µL of 10 mg/mL hexacosane (Millipore Sigma, ≥98.0% (GC)) in hexane (Sigma Aldrich, 1043671000, ACS reagent, EMSURE®) as an internal standard. The vials were incubated in a room temperature shaker for 15 minutes. Then, the liquid was removed and placed into a new 2 mL amber vial with 150 µL insert and sealed immediately with Teflon tape wrapped around the base of the cap. Three replicates were prepared for each genotype. Samples were shipped overnight on dry ice to Bucknell University where they were stored at-80°C before analysis with gas chromatography/mass spectrometry (GC-MS). Measurements were performed on an Agilent 8860/5977B GC-MS with an HP-5MS-UI column (30 m long, 0.25 mm inner diameter, 0.25 µm film thickness) using ultra-high purity helium as the carrier gas. Liquid samples (1 µL) were injected into a 280°C splitless inlet using an autosampler fitted with a 10 µL syringe (MPS TD/robotic; Gerstel GmbH). The oven started at 40°C with 1 min hold, then ramped to 280°C at 10°C/min, with a final 280°C hold for 10 min. Compound identification was performed using mass spectral interpretation coupled with retention indices that were calibrated against a mixture of normal alkanes (Restek; P/N 31080). All samples were injected 3 times in immediate succession, in numerical order. Data was processed with Agilent MassHunter Quantitative Analysis 10.0 for relative quantification using the ion signal with the largest intensity in the spectrum as the quantifier; the next two largest ion signals were used as qualifiers. Peak areas for all compounds were normalized to hexacosane as the internal standard.

### High Sugar Diet

1 to 2-day old males were put on high-sugar diet (0.75 M Sucrose) for 10 days upon eclosion and flipped onto new vials of food every day. Sleep and sensitivity to HO was assessed as previously described. Experiment was performed with normal Archon media.

### Survival curves

Survival curves for starvation assays and hyperoxia assays are all plotted as Kaplan-Meier graphs. Log-rank analysis was performed using GraphPad Prism. All experiments were performed with a minimum of three independent trials and yielded statistically similar results, except where noted. Graphs and *p*-values in figures are from representative trials.

## Statistical analysis

We assessed the normality of our data using the D’Agostino-Pearson omnibus normality test and the Shapiro-Wilk normality test, which have good power properties over a wide range of distributions.^71^ For datasets that passed both normality tests, we used the unpaired Student’s t-test with Welch’s correction when comparing two groups and the one-way ANOVA (Dunn’s multiple comparisons) when comparing three or more groups. For data sets that failed either one of the normality tests, we used the Mann-Whitney U test when comparing two groups and the Kruskal-Wallis test with Dunn’s post hoc test when comparing three or more groups. Significance is expressed as p values (n.s., p > 0.05; *, p < 0.05; **, p < 0.01; *** p < 0.001; **** p < 0.0001).

## AUTHOR CONTRIBUTIONS

SJT, CEK, and MSH conceived the experiments. Experiments were performed and analyzed by SJT (husbandry, hyperoxia survival, DAM sleep, RNA-sequencing sample collection and secondary data analysis, antibiotic feeding, starvation survival, hyperoxia and starvation combined survival, TAG measurement, respirometry, metabolomics sample collection and secondary data analysis), CEK (husbandry, Sleep TAG measurements, hyperoxia and starvation combined survival, Desiccation/Dehydration, *Bmm* and HSD Sleep and HO survival), JL (husbandry, DAM sleep normoxia vs hyperoxia, hyperoxia and starvation combined survival, respirometry, metabolomics secondary data analysis), KO (husbandry, DAM sleep normoxia vs hyperoxia, starvation survival, TAG measurement), JAG (Sleep TAG measurement, respirometry), TYC (RNA-sequencing secondary data analysis), CL (*Bmm* and HSD TAG measurement, respirometry), RS (RNA-sequencing secondary data analysis), MSJ (RNA-sequencing secondary data analysis), SJP (feeding), JMH (RNA-sequencing secondary data analysis), MU (gut dissections and DHE staining), WWJ (feeding), DBC (CHC GC/MS and analysis). SJT, CEK, JCC, and MSH made intellectual contributions. SJT, CEK, JCC, and MSH designed the figures and wrote the manuscript.

## Supporting information

RNA seq

metabolomics

## ACKNOWLEDGMENTS

We thank all members of the Shirasu-Hiza and Canman labs for support, discussions, and feedback. The RNA-sequencing was funded in part through the NIH/NCI CancerCenter Support Grant P30CA013696 and used the Genomics and High Throughput Screening Shared Resource. We also thank Daniel Bushey, Nicholas Stavropoulos, Rob Jackson, and Amita Sehgal for fly lines. Work was supported by NIH F31AG074664 (SJT), GSAS-Leadership Alliance and Amgen Scholars programs (JL), Columbia University SURF (KO), NIH F31AG079601 (JAG), NSF GRFP (TYC), NSF GRFP (CL), NIH-NIGMS T32 Fellowship GM067545 (MSJ), NIH-NIGMS MIRA R35GM128687 (JMH), NSF CAREER award 2045673 (JMH), AFAR Glenn Foundation Postdoctoral Fellowship for Aging Research (MU), NIH R01GM117407 (JCC), NIH R01GM130764 (JCC), NIH R01DC020031 (WWJ), NSF IOS-2035286 (DBC), NIH R35GM127049 (MSH) and NIH R01AG045842 (MSH).

## DATA AVAILABILITY

The authors declare that all data supporting the findings of this study are available, including replicate experiments, and will be made available upon reasonable request to the corresponding author, Dr. Mimi Shirasu-Hiza.

## CONFLICT OF INTEREST

The authors declare no competing interests.

## Notes

### Competing Interest Statement

The authors have declared no competing interest.

## REFERENCES

1 Campbell, S. S. & Tobler, I. Animal sleep: a review of sleep duration across phylogeny. Neurosci Biobehav Rev 8, 269–300 (1984). 10.1016/0149-7634(84)90054-x

2 Cirelli, C. & Tononi, G. Is sleep essential? PLoS Biol 6, e216 (2008). 10.1371/journal.pbio.0060216

3 Miyazaki, S., Liu, C.-Y. & Hayashi, Y. Sleep in vertebrate and invertebrate animals, and insights into the function and evolution of sleep. Neuroscience Research 118, 3–12 (2017). 10.1016/j.neures.2017.04.017

4 Turrens, J. F. Mitochondrial formation of reactive oxygen species. J Physiol 552, 335–344 (2003). 10.1113/jphysiol.2003.049478

5 Keene, A. C. & Duboue, E. R. The origins and evolution of sleep. J Exp Biol 221 (2018). https://doi.org:ARTN jeb159533 10.1242/jeb.159533

6 Nedeltcheva, A. V. & Scheer, F. A. Metabolic effects of sleep disruption, links to obesity and diabetes. Curr Opin Endocrinol Diabetes Obes 21, 293–298 (2014). 10.1097/MED.0000000000000082

7 Taheri, S. The link between short sleep duration and obesity: we should recommend more sleep to prevent obesity. Archives of Disease in Childhood 91, 881–884 (2006). 10.1136/adc.2005.093013

8 Brum, M. C. B. et al. Night shift work, short sleep and obesity. Diabetol Metab Syndr 12, 13 (2020). 10.1186/s13098-020-0524-9

9 Knutson, K. L., Ryden, A. M., Mander, B. A. & Van Cauter, E. Role of sleep duration and quality in the risk and severity of type 2 diabetes mellitus. Arch Intern Med 166, 1768–1774 (2006). 10.1001/archinte.166.16.1768

10 Cappuccio, F. P., Cooper, D., D’Elia, L., Strazzullo, P. & Miller, M. A. Sleep duration predicts cardiovascular outcomes: a systematic review and meta-analysis of prospective studies. Eur Heart J 32, 1484–1492 (2011). 10.1093/eurheartj/ehr007

11 Lim, A. S., Kowgier, M., Yu, L., Buchman, A. S. & Bennett, D. A. Sleep Fragmentation and the Risk of Incident Alzheimer’s Disease and Cognitive Decline in Older Persons. Sleep 36, 1027–1032 (2013). 10.5665/sleep.2802

12 Musiek, E. S., Xiong, D. D. & Holtzman, D. M. Sleep, circadian rhythms, and the pathogenesis of Alzheimer disease. Exp Mol Med 47, e148 (2015). 10.1038/emm.2014.121

13 Sterniczuk, R., Theou, O., Rusak, B. & Rockwood, K. Sleep disturbance is associated with incident dementia and mortality. Curr Alzheimer Res 10, 767–775 (2013). 10.2174/15672050113109990134

14 Holth, J. K. et al. The sleep-wake cycle regulates brain interstitial fluid tau in mice and CSF tau in humans. Science 363, 880–884 (2019). 10.1126/science.aav2546

15 Winer, J. R. et al. Sleep Disturbance Forecasts beta-Amyloid Accumulation across Subsequent Years. Curr Biol 30, 4291–4298 e4293 (2020). 10.1016/j.cub.2020.08.017

16 Schenck, C. H., Boeve, B. F. & Mahowald, M. W. Delayed emergence of a parkinsonian disorder or dementia in 81% of older men initially diagnosed with idiopathic rapid eye movement sleep behavior disorder: a 16-year update on a previously reported series. Sleep Med 14, 744–748 (2013). 10.1016/j.sleep.2012.10.009

17 Bo, Y. et al. Sleep and the Risk of Chronic Kidney Disease: A Cohort Study. J Clin Sleep Med 15, 393–400 (2019). 10.5664/jcsm.7660

18 Wijarnpreecha, K., Thongprayoon, C., Panjawatanan, P. & Ungprasert, P. Short sleep duration and risk of nonalcoholic fatty liver disease: A systematic review and meta-analysis. J Gastroenterol Hepatol 31, 1802–1807 (2016). 10.1111/jgh.13391

19 Kim, C.-W. et al. Sleep duration and quality in relation to non-alcoholic fatty liver disease in middle-aged workers and their spouses. Journal of Hepatology 59, 351–357 (2013). 10.1016/j.jhep.2013.03.035

20 Orr, W. C., Fass, R., Sundaram, S. S. & Scheimann, A. O. The effect of sleep on gastrointestinal functioning in common digestive diseases. Lancet Gastroenterol Hepatol 5, 616–624 (2020). 10.1016/S2468-1253(19)30412-1

21 Besedovsky, L., Lange, T. & Born, J. Sleep and immune function. Pflugers Arch 463, 121–137 (2012). 10.1007/s00424-011-1044-0

22 Hublin, C., Partinen, M., Koskenvuo, M. & Kaprio, J. Sleep and mortality: a population-based 22-year follow-up study. Sleep 30, 1245–1253 (2007). 10.1093/sleep/30.10.1245

23 Mazzotti, D. R. et al. Human longevity is associated with regular sleep patterns, maintenance of slow wave sleep, and favorable lipid profile. Frontiers in Aging Neuroscience 6 (2014). 10.3389/fnagi.2014.00134

24 Gallicchio, L. & Kalesan, B. Sleep duration and mortality: a systematic review and meta-analysis. J Sleep Res 18, 148–158 (2009). 10.1111/j.1365-2869.2008.00732.x

25 Shaw, P. J., Cirelli, C., Greenspan, R. J. & Tononi, G. Correlates of sleep and waking in Drosophila melanogaster. Science 287, 1834–1837 (2000). 10.1126/science.287.5459.1834

26 Hendricks, J. C. et al. Rest in Drosophila is a sleep-like state. Neuron 25, 129–138 (2000). 10.1016/s0896-6273(00)80877-6

27 Hill, V. M. et al. A bidirectional relationship between sleep and oxidative stress in Drosophila. PLOS Biology 16, e2005206 (2018). 10.1371/journal.pbio.2005206

28 Reimund, E. The free radical flux theory of sleep. Med Hypotheses 43, 231–233 (1994). 10.1016/0306-9877(94)90071-x

29 D’Almeida, V. et al. Sleep deprivation induces brain region-specific decreases in glutathione levels. Neuroreport 9, 2853–2856 (1998). 10.1097/00001756-199808240-00031

30 Eiland, M. M. et al. Increases in amino-cupric-silver staining of the supraoptic nucleus after sleep deprivation. Brain Res 945, 1–8 (2002). 10.1016/s0006-8993(02)02448-4

31 Ramanathan, L., Gulyani, S., Nienhuis, R. & Siegel, J. M. Sleep deprivation decreases superoxide dismutase activity in rat hippocampus and brainstem. Neuroreport 13, 1387–1390 (2002). 10.1097/00001756-200208070-00007

32 Silva, R. H. et al. Role of hippocampal oxidative stress in memory deficits induced by sleep deprivation in mice. Neuropharmacology 46, 895–903 (2004). 10.1016/j.neuropharm.2003.11.032

33 Everson, C. A., Laatsch, C. D. & Hogg, N. Antioxidant defense responses to sleep loss and sleep recovery. Am J Physiol Regul Integr Comp Physiol 288, R374–383 (2005). 10.1152/ajpregu.00565.2004

34 Singh, R., Kiloung, J., Singh, S. & Sharma, D. Effect of paradoxical sleep deprivation on oxidative stress parameters in brain regions of adult and old rats. Biogerontology 9, 153–162 (2008). 10.1007/s10522-008-9124-z

35 Villafuerte, G. et al. Sleep deprivation and oxidative stress in animal models: a systematic review. Oxid Med Cell Longev 2015, 234952 (2015). 10.1155/2015/234952

36 Bushey, D., Huber, R., Tononi, G. & Cirelli, C. Drosophila mutants have reduced sleep and impaired memory. J Neurosci 27, 5384–5393 (2007). 10.1523/Jneurosci.0108-07.2007

37 Stavropoulos, N. & Young, M. W. insomniac and Cullin-3 regulate sleep and wakefulness in Drosophila. Neuron 72, 964–976 (2011). 10.1016/j.neuron.2011.12.003

38 Kume, K., Kume, S., Park, S. K., Hirsh, J. & Jackson, F. R. Dopamine is a regulator of arousal in the fruit fly. J Neurosci 25, 7377–7384 (2005). 10.1523/Jneurosci.2048-05.2005

39 Shi, M., Yue, Z. F., Kuryatov, A., Lindstrom, J. M. & Sehgal, A. Identification of Redeye, a new sleep-regulating protein whose expression is modulated by sleep amount. Elife 3 (2014). https://doi.org:ARTN e01473 10.7554/eLife.01473

40 Pfeiffenberger, C. & Allada, R. Cul3 and the BTB adaptor insomniac are key regulators of sleep homeostasis and a dopamine arousal pathway in Drosophila. PLoS Genet 8, e1003003 (2012). 10.1371/journal.pgen.1003003

41 Kempf, A., Song, S. M., Talbot, C. B. & Miesenbock, G. A potassium channel beta-subunit couples mitochondrial electron transport to sleep. Nature 568, 230–234 (2019). 10.1038/s41586-019-1034-5

42 Vaccaro, A. et al. Sleep Loss Can Cause Death through Accumulation of Reactive Oxygen Species in the Gut. Cell 181, 1307-+ (2020). 10.1016/j.cell.2020.04.049

43 Bonnet, M. H. & Arand, D. L. We are chronically sleep deprived. Sleep 18, 908–911 (1995). 10.1093/sleep/18.10.908

44 Chattu, V. K. et al. The Global Problem of Insufficient Sleep and Its Serious Public Health Implications. Healthcare (Basel) 7 (2018). 10.3390/healthcare7010001

45 Wickwire, E. M., Geiger-Brown, J., Scharf, S. M. & Drake, C. L. Shift Work and Shift Work Sleep Disorder: Clinical and Organizational Perspectives. Chest 151, 1156–1172 (2017). 10.1016/j.chest.2016.12.007

46 Ohayon, M. M. Epidemiological Overview of sleep Disorders in the General Population. Sleep Med Res 2, 1–9 (2011). 10.17241/smr.2011.2.1.1

47 Hill, V. M. et al. A bidirectional relationship between sleep and oxidative stress in Drosophila. Plos Biology 16 (2018). https://doi.org:ARTN 2005206 10.1371/journal.pbio.2005206

48 Pfeiffenberger, C. & Allada, R. Cul3 and the BTB Adaptor Are Key Regulators of Sleep Homeostasis and a Dopamine Arousal Pathway in Drosophila. Plos Genet 8 (2012). https://doi.org:ARTN e100300310.1371/journal.pgen.1003003

49 Tener, S. J. et al. Neuronal knockdown of Cullin3 as a Drosophila model of autism spectrum disorder. Sci Rep 14, 1541 (2024). 10.1038/s41598-024-51657-9

50 Gilestro, G. F. & Cirelli, C. pySolo: a complete suite for sleep analysis in Drosophila. Bioinformatics 25, 1466–1467 (2009). 10.1093/bioinformatics/btp237

51 Sykiotis, G. P. & Bohmann, D. Keap1/Nrf2 signaling regulates oxidative stress tolerance and lifespan in Drosophila. Dev Cell 14, 76–85 (2008). 10.1016/j.devcel.2007.12.002

52 Owen, J. B. & Butterfield, D. A. Measurement of oxidized/reduced glutathione ratio. Methods Mol Biol 648, 269–277 (2010). 10.1007/978-1-60761-756-3_18

53 Douglas, A. E. The Drosophila model for microbiome research. Lab Anim (NY) 47, 157–164 (2018). 10.1038/s41684-018-0065-0

54 Kunst, C. et al. The Influence of Gut Microbiota on Oxidative Stress and the Immune System. Biomedicines 11 (2023). 10.3390/biomedicines11051388

55 Quintero, P., Gonzalez-Muniesa, P., Garcia-Diaz, D. F. & Martinez, J. A. Effects of hyperoxia exposure on metabolic markers and gene expression in 3T3-L1 adipocytes. J Physiol Biochem 68, 663–669 (2012). 10.1007/s13105-012-0169-8

56 Stellingwerff, T. et al. Effects of hyperoxia on skeletal muscle carbohydrate metabolism during transient and steady-state exercise. J Appl Physiol (1985) 98, 250–256 (2005). 10.1152/japplphysiol.00897.2004

57 Skandalis, D. A., Stuart, J. A. & Tattersall, G. J. Responses of Drosophila melanogaster to atypical oxygen atmospheres. J Insect Physiol 57, 444–451 (2011). 10.1016/j.jinsphys.2011.01.005

58 Shukla, A. K. et al. Metabolomic Analysis Provides Insights on Paraquat-Induced Parkinson-Like Symptoms in Drosophila melanogaster. Mol Neurobiol 53, 254–269 (2016). 10.1007/s12035-014-9003-3

59 Lin, J., Handschin, C. & Spiegelman, B. M. Metabolic control through the PGC-1 family of transcription coactivators. Cell Metab 1, 361–370 (2005). 10.1016/j.cmet.2005.05.004

60 Nowak, N. et al. Rapid and reversible control of human metabolism by individual sleep states. Cell Rep 37, 109903 (2021). 10.1016/j.celrep.2021.109903

61 Zhang, S. et al. Metabolic flexibility during sleep. Sci Rep 11, 17849 (2021). 10.1038/s41598-021-97301-8

62 Kayaba, M. et al. Energy metabolism differs between sleep stages and begins to increase prior to awakening. Metabolism 69, 14–23 (2017). 10.1016/j.metabol.2016.12.016

63 Wang, Z. et al. Desiccation resistance differences in Drosophila species can be largely explained by variations in cuticular hydrocarbons. Elife 11 (2022). 10.7554/eLife.80859

64 Bedont, J. L. et al. Chronic sleep loss sensitizes Drosophila melanogaster to nitrogen stress. Current Biology 33, 1613-+ (2023). 10.1016/j.cub.2023.03.008

65 Wang, L. et al. Spatially resolved isotope tracing reveals tissue metabolic activity. Nat Methods 19, 223–230 (2022). 10.1038/s41592-021-01378-y

66 Wilinski, D. et al. Rapid metabolic shifts occur during the transition between hunger and satiety in Drosophila melanogaster. Nat Commun 10 (2019). https://doi.org:ARTN 405210.1038/s41467-019-11933-z

67 Kierans, S. J. & Taylor, C. T. Regulation of glycolysis by the hypoxia-inducible factor (HIF): implications for cellular physiology. J Physiol 599, 23–37 (2021). 10.1113/JP280572

68 Schindelin, J., et al. Fiji: an open-source platform for biological-image analysis. Nat Methods 9, 676–682 (2012). 10.1038/nmeth.2019

69 Deshpande, S. A. et al. Quantifying Drosophila food intake: comparative analysis of current methodology. Nat Methods 11, 535–540 (2014). 10.1038/nmeth.2899

70 Yatsenko, A. S., Marrone, A. K., Kucherenko, M. M. & Shcherbata, H. R. Measurement of metabolic rate in Drosophila using respirometry. J Vis Exp, e51681 (2014). 10.3791/51681

71 Yap, B. W. & Sim, C. H. Comparisons of various types of normality tests. J Stat Comput Sim 81, 2141–2155 (2011). 10.1080/00949655.2010.520163

