## supplemental figures for "Investigating the consequences of chronic short sleep for metabolism and survival of oxidative stress"

Tener et al., Figure S1

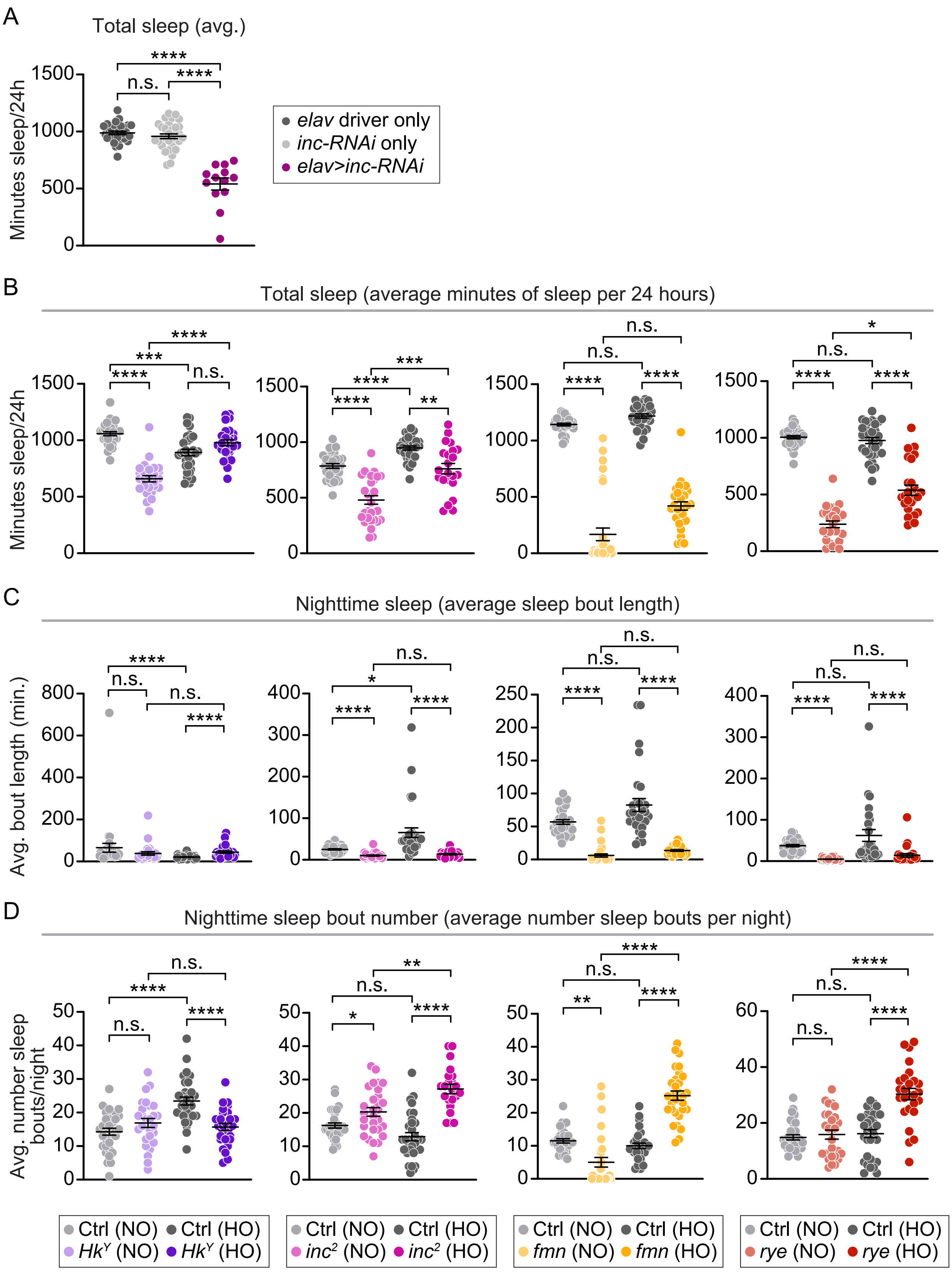

Tener et al., Figure S2

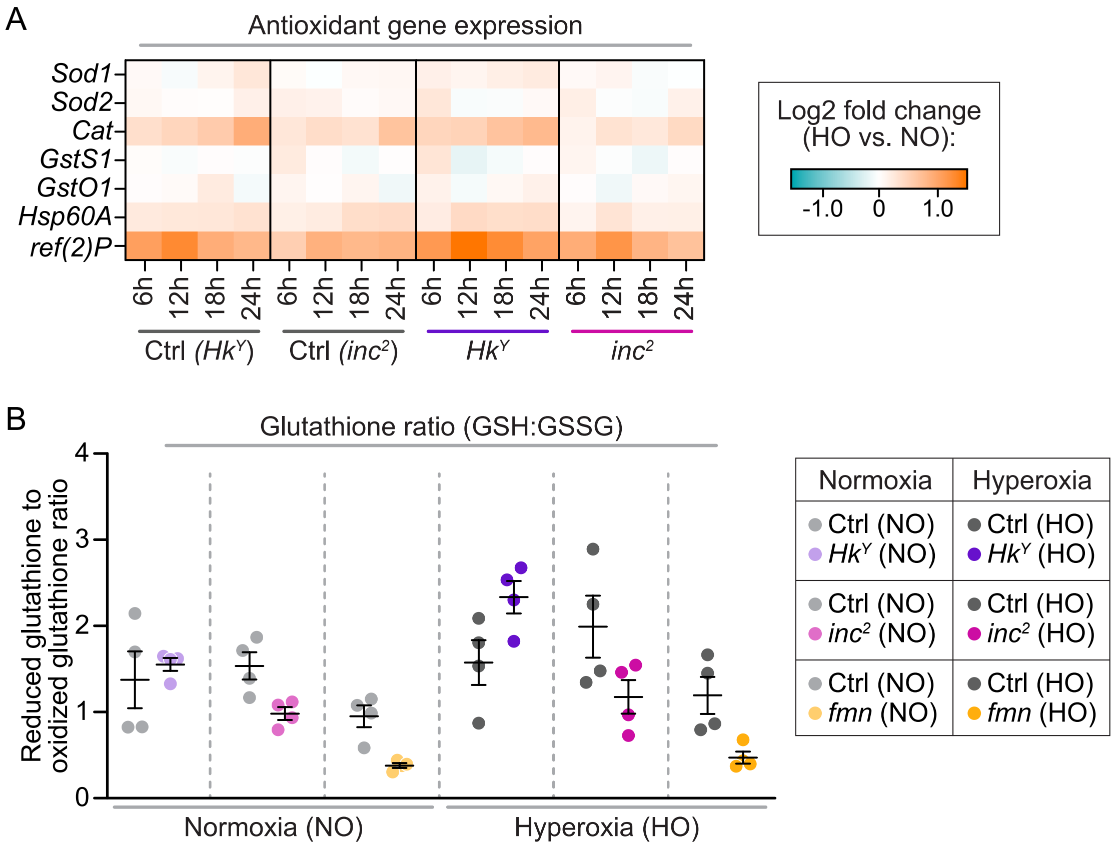

Tener et al., Figure S3

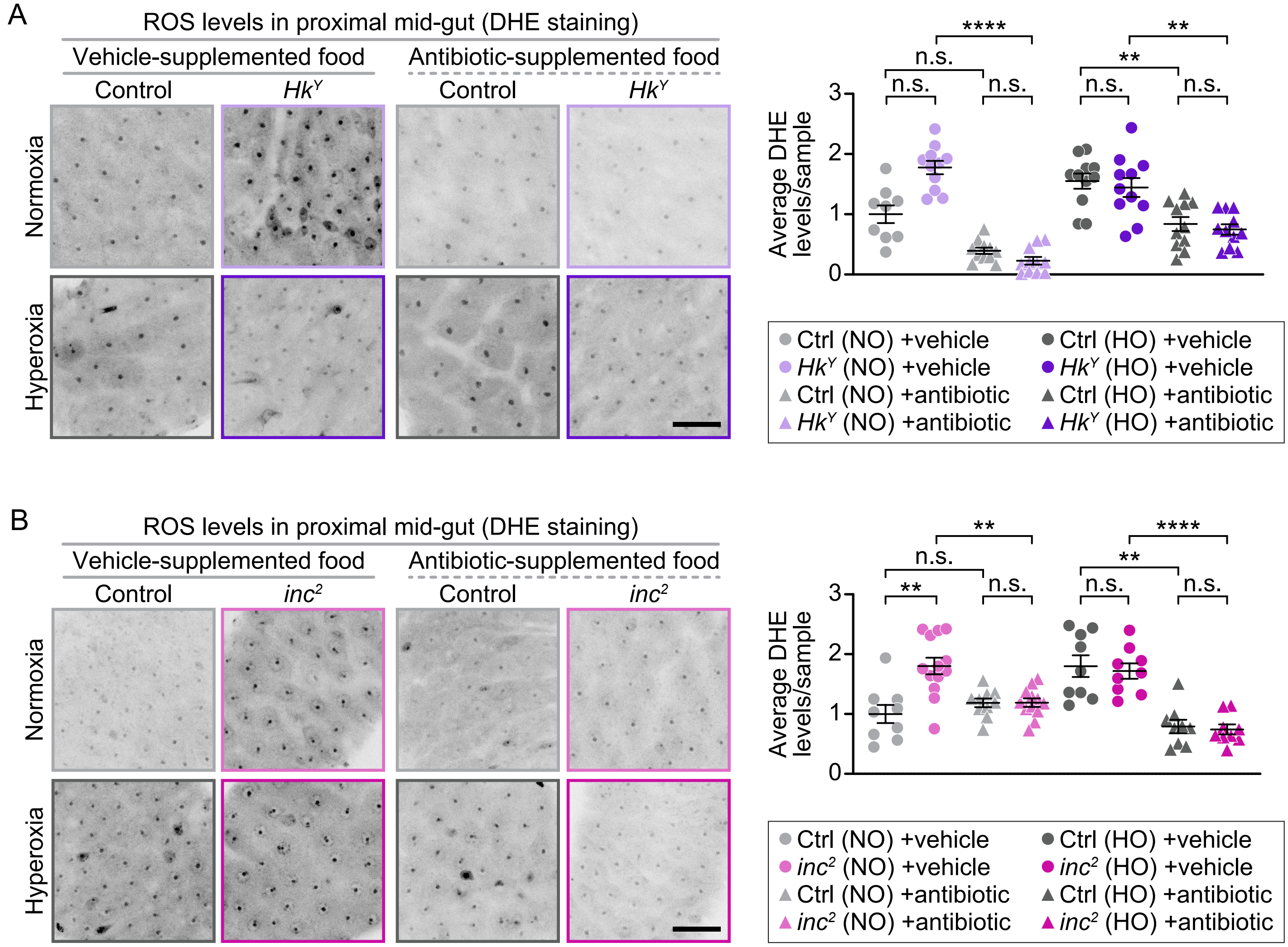

Tener et al., Figure S4

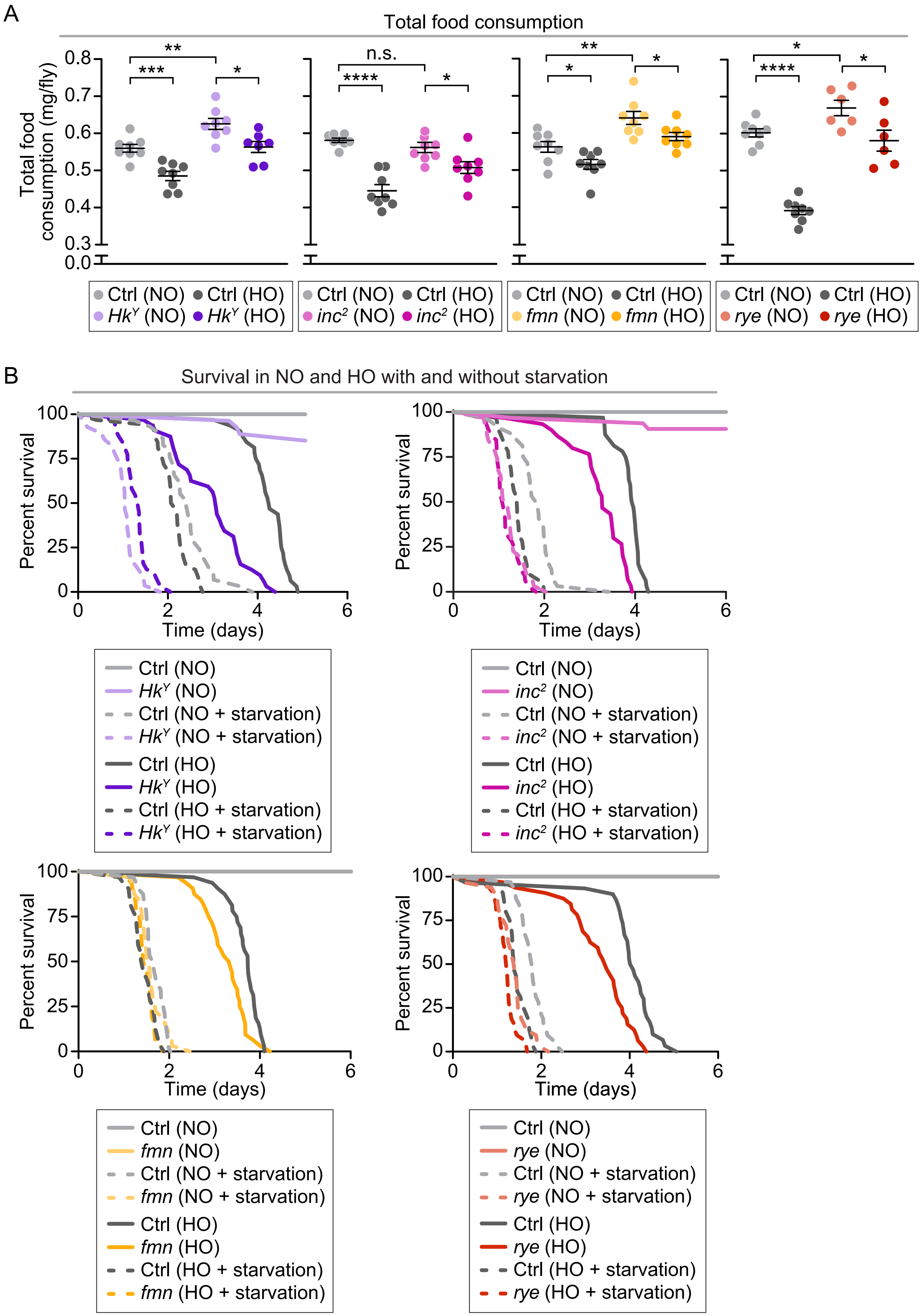

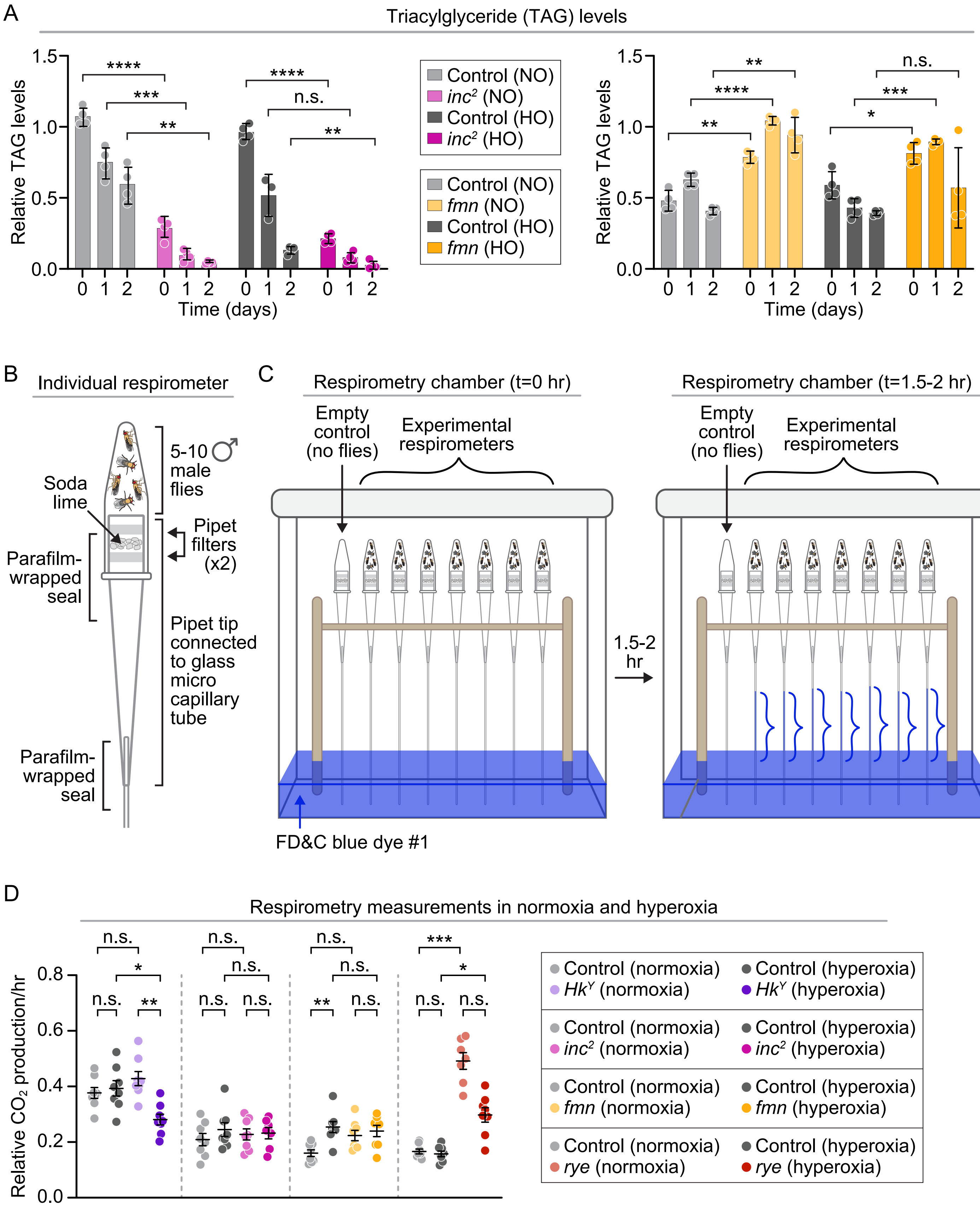
A
Triacylglyceride (TAG) levels
Relative TAG levels
1.5
1.0
0.5
0.0
****
***
**
****
n.s.
**
0 1 2 0 1 2 0 1 2 0 1 2
Time (days)
Control (NO)
*$inc^2$* (NO)
Control (HO)
*$inc^2$* (HO)
Control (NO)
*fmn* (NO)
Control (HO)
*fmn* (HO)
**
****
**
n.s.
***
*
B
Individual respirometer
Soda lime
5-10 ♂ male flies
Pipet filters (x2)
Parafilm-wrapped seal
Pipet tip connected to glass micro capillary tube
Parafilm-wrapped seal
C
Respirometry chamber (t=0 hr)
Empty control (no flies)
Experimental respirometers
FD&C blue dye #1
1.5-2 hr
Respirometry chamber (t=1.5-2 hr)
Empty control (no flies)
Experimental respirometers
D
Respirometry measurements in normoxia and hyperoxia
Relative $CO_2$ production/hr
0.8
0.6
0.4
0.2
0.0
n.s.
*
n.s.
**
n.s.
n.s.
n.s.
n.s.
n.s.
n.s.
**
n.s.
***
*
n.s.
n.s.
Control (normoxia)
*$Hk^Y$* (normoxia)
Control (hyperoxia)
*$Hk^Y$* (hyperoxia)
Control (normoxia)
*$inc^2$* (normoxia)
Control (hyperoxia)
*$inc^2$* (hyperoxia)
Control (normoxia)
*fmn* (normoxia)
Control (hyperoxia)
*fmn* (hyperoxia)
Control (normoxia)
*rye* (normoxia)
Control (hyperoxia)
*rye* (hyperoxia)

Tener et al., Figure S6

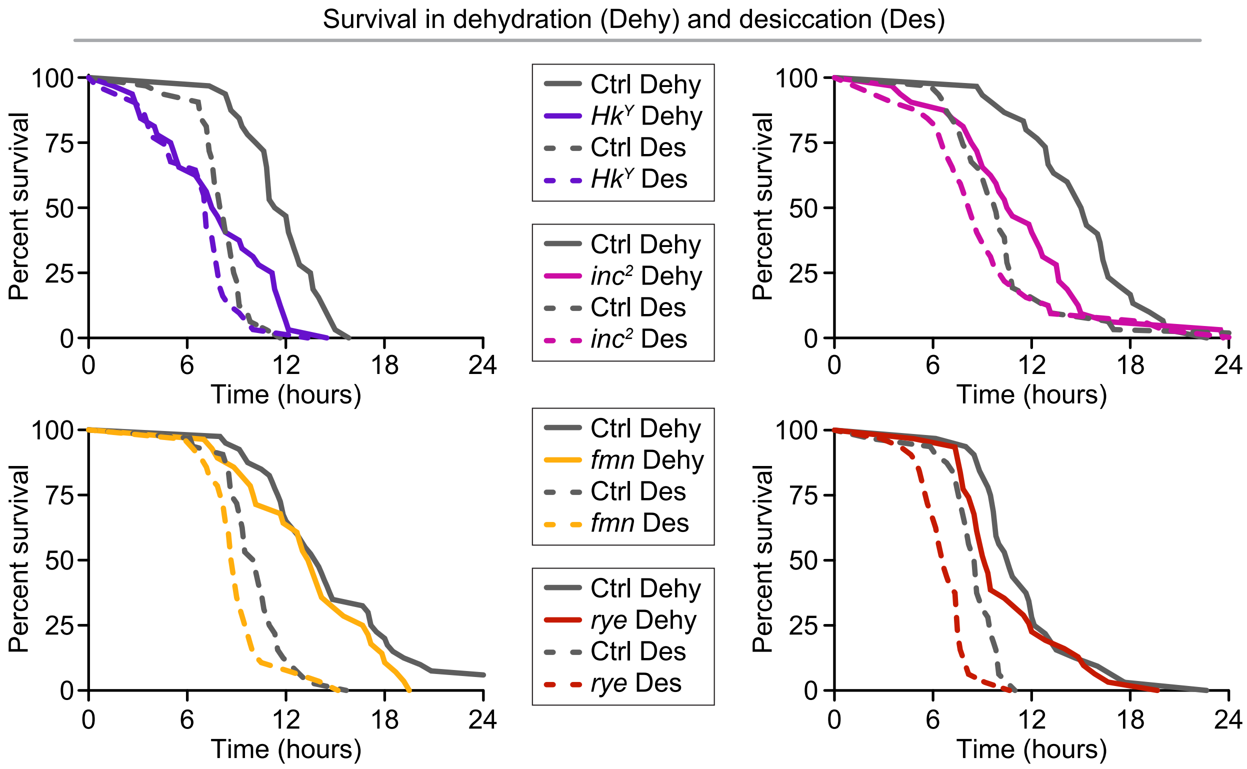
